# From proteins to polysaccharides: lifestyle and genetic evolution of *Coprothermobacter proteolyticus*

**DOI:** 10.1101/280602

**Authors:** B. J. Kunath, F. Delogu, A.E. Naas, M.Ø. Arntzen, V.G.H. Eijsink, B. Henrissat, T.R. Hvidsten, P.B. Pope

## Abstract

Microbial communities that degrade lignocellulosic biomass are typified by high levels of species- and strain-level complexity as well as synergistic interactions between both cellulolytic and non-cellulolytic microorganisms. *Coprothermobacter proteolyticus* frequently dominates thermophilic, lignocellulose-degrading communities with wide geographical distribution, which is in contrast to reports that it ferments proteinaceous substrates and is incapable of polysaccharide hydrolysis. Here we deconvolute a highly efficient cellulose-degrading consortium (SEM1b) that is co-dominated by *Clostridium (Ruminiclostridium) thermocellum*- and multiple heterogenic strains affiliated to *C. proteolyticus*. Metagenomic analysis of SEM1b recovered metagenome-assembled genomes (MAGs) for each constituent population, whilst in parallel two novel strains of *C. proteolyticus* were successfully isolated and sequenced. Annotation of all *C. proteolyticus* genotypes (two strains and one MAG) revealed their genetic acquisition of carbohydrate-active enzymes (CAZymes), presumably derived from horizontal gene transfer (HGT) events involving polysaccharide-degrading Firmicutes or Thermotogae-affiliated populations that are historically co-located. HGT material included a saccharolytic operon, from which a CAZyme was biochemically characterized and demonstrated hydrolysis of multiple hemicellulose polysaccharides. Finally, temporal genome-resolved metatranscriptomic analysis of SEM1b revealed expression of *C. proteolyticus* CAZymes at different SEM1b life-stages as well as co-expression of CAZymes from multiple SEM1b populations, inferring deeper microbial interactions that are dedicated towards community degradation of cellulose and hemicellulose. We show that *C. proteolyticus*, a ubiquitous keystone population, consists of closely related strains that have adapted via HGT to presumably degrade both oligo- and longer polysaccharides present in decaying plants and microbial cell walls, thus explaining its dominance in thermophilic anaerobic digesters on a global scale.

## INTRODUCTION

The anaerobic digestion of plant biomass profoundly shapes innumerable ecosystems, ranging from the gastrointestinal tracts of humans and other mammals to those that drive industrial applications such as biofuel generation. Biogas reactors are one of the most commonly studied anaerobic systems, yet many keystone microbial populations and their metabolic processes are poorly understood due to a lack of cultured or genome sampled representatives. *Coprothermobacter* spp. are frequently observed in high abundance in thermophilic anaerobic systems, where they are believed to exert strong protease activity whilst generating hydrogen and acetate, key intermediate metabolites for biogas production (Tandishabo et al 2012). Molecular techniques have shown that their levels range from 10 to 90% of the total microbial community, irrespective of bioreactors being operated on lignocellulose- or protein-rich substrates (**Figure 1**). Despite their promiscuous distribution, global abundance and key role in biogas production, only two species have been described: *Coprothermobacter platensis* (Etchebehere et al 1998) and *Coprothermobacter proteolyticus* (Ollivier et al 1985). These two species and their inherent phenotypes have formed the predictive basis for the majority of *Coprothermobacter*-dominated systems described to date. Recent studies have illustrated that *C. proteolyticus* populations in anaerobic biogas reactors form cosmopolitan assemblages of closely related strains that are hitherto unresolved (Hagen et al 2017).

**Figure 1.**
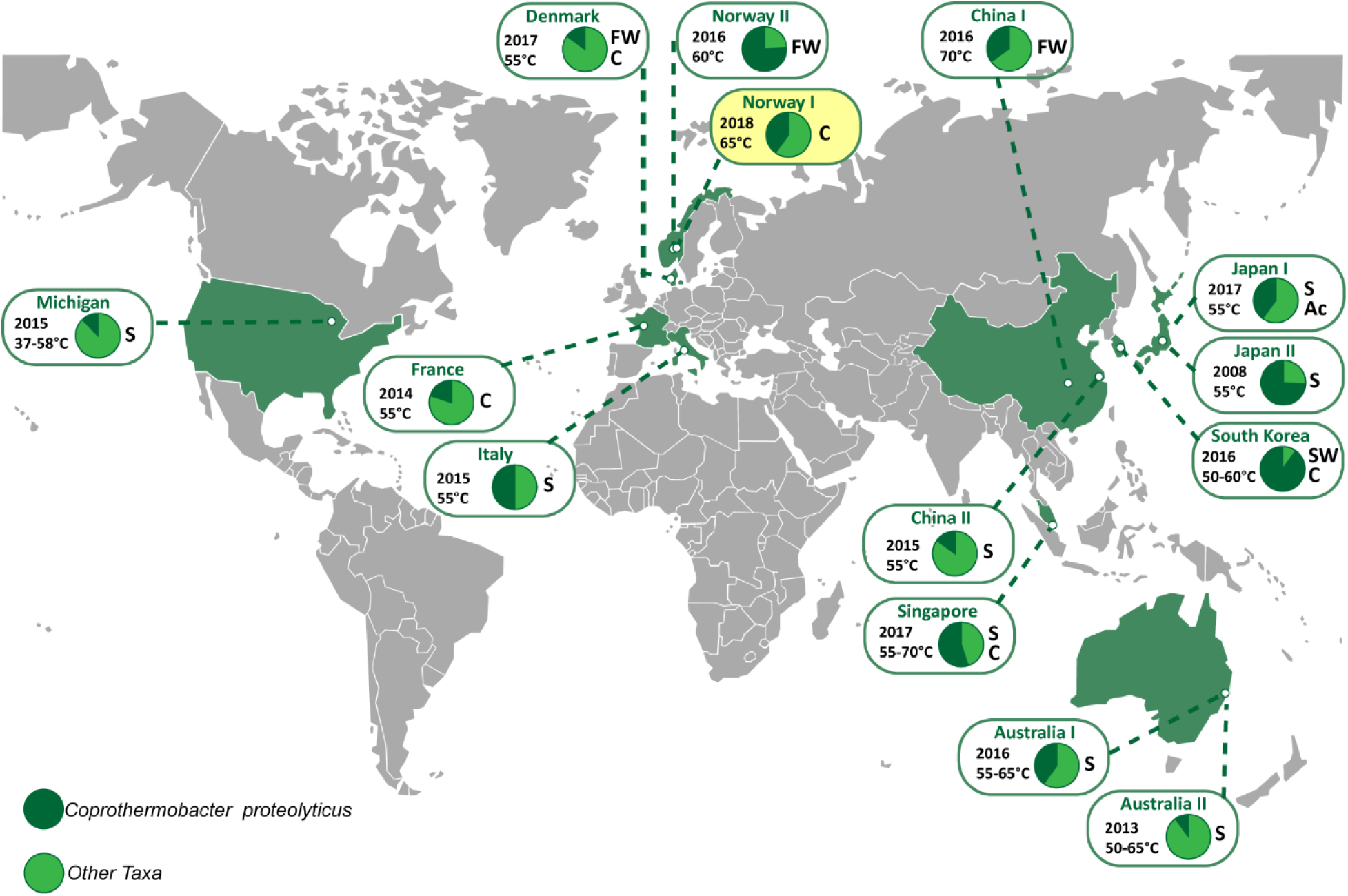
Global distribution of *Coprothermobacter* proteolyticus-affiliated populations in anaerobic biogas reactors. Charts indicate relative 16S rRNA gene abundance of OTUs affiliated to *C. proteolyticus* (dark green), in comparison to the total community (light green). The year of publication, reactor temperature and substrate (C: cellulose, FW: food waste, S: sludge, SW: Seaweed, Ac: acetate) is indicated (details in **Table S1**). The SEM1b consortium analyzed in this study is highlighted in yellow.

Frequently in nature, microbial populations are composed of multiple strains with genetic heterogeneity (Kashtan et al 2014, Schloissnig et al 2013). Studies of strain-level populations have been predominately performed with the human microbiome and especially the gut microbiota (Bron et al 2012, Spanogiannopoulos et al 2016). The reasons for strain diversification and their coexistence remain largely unknown (Ellegaard and Engel 2016), however several mechanisms have been hypothesized, such as: micro-niche selection (Hunt et al 2008, Kashtan et al 2014), host selection (McLoughlin et al 2016), cross-feed interactions (Rosenzweig et al 1994, Zelezniak et al 2015) and phage selection (Rodriguez-Valera et al 2009). Studies of isolated strains have shown that isolates can differ in a multitude of ways, including virulence and drug resistance (Gill et al 2005, Sharon et al 2013, Solheim et al 2009), motility (Zunino et al 1994) and nutrient utilization (Siezen et al 2010). Strain-level genomic variations typically consist of single-nucleotide variants (SNVs) as well as acquisition/loss of genomic elements such as genes, operons or plasmids via horizontal gene transfer (HGT) (Koskella and Vos 2015, Tettelin et al 2005, Treangen and Rocha 2011). Variability in gene content caused by HGT is typically attributed to phage-related genes and other genes of unknown function (Ochman et al 2000), and can give rise to ecological adaptation, niche differentiation and eventually speciation (Bendall et al 2016, Biller et al 2015, Shapiro et al 2012). Although differences in genomic features can be accurately characterized in isolated strains, it has been difficult to capture such information using culture-independent approaches such as metagenomics. Advances in bioinformatics have improved taxonomic profiling of microbial communities from phylum to species level but it remains difficult to profile similar strains from metagenomes and compare them with the same level of resolution obtained by comparison of isolate genomes (Truong et al 2017). Since closely-related strains can also differ in gene expression (González-Torres et al 2015), being able to distinguish the expression profiles of individual strains in a broader ecological context is elemental to understanding the influence they exert towards the overall community function.

In this study, a novel population of *C. proteolyticus* that included multiple closely related strains, was observed within a simplistic biogas-producing consortium enriched on cellulose (hereafter referred to as SEM1b). Using a combined metagenomic and culture-dependent approach, two strains and a metagenome-assembled genome (MAG) affiliated to *C. proteolyticus* were recovered and genetically compared to the only available type strain, *C. proteolyticus* DSM 5265 (Alexiev et al 2014). Notable genomic differences included the acquisition of an operon (region-A) encoding carbohydrate-active enzymes (CAZymes), which inferred that *C. proteolyticus* has adapted to take advantage of longer polysaccharides. Enzymology was used to further support our hypothesis that the CAZymes within region-A are functionally active. We further examined the saccharolytic potential of our recovered *C. proteolyticus* population in a broader community context, by examining genome-resolved temporal metatranscriptomic data generated from the SEM1b consortium. Collective analysis highlighted the time-specific polysaccharide-degrading activity that *C. proteolyticus* exerts in a cellulolytic microbial community.

## MATERIALS AND METHODS

### Origin of samples and generation of the SEM1b consortium

An inoculum (100μl) was collected from a lab-scale biogas reactor (Reactor TD) fed with manure and food waste and run at 55°C. The TD reactor originated itself from a thermophilic (60°C) biogas plant (Frevar) fed with food waste and manure in Fredrikstad, Norway. Our research groups have previously studied the microbial communities in both the Frevar plant (Hagen et al 2017) and the TD bioreactor (Zamanzadeh et al 2016), which provided a detailed understanding of the original microbial community. The inoculum was transferred for serial dilution and enrichment to an anaerobic serum bottle and containing the rich ATCC medium 1943, with cellobiose substituted for 10g/L of cellulose in the form of Borregaard Advanced Lignin technology (BALI^TM^) treated Norway spruce (Rødsrud et al 2012). Our enrichment was incubated at 65°C with the lesser objective to study community biomass conversion at the upper temperature limits of methanogensis. After an initial growth cycle, an aliquot was removed and used for a serial dilution to extinction experiment. Briefly, a 100μl sample was transferred to a new 100ml bottle containing 60ml of anaerobic medium, mixed and 100μl was directly transferred again to a new one (six serial transfers in total). The consortium at maximum dilution that retained the cellulose-degrading capability (SEM1b) was retained for the present work, and aliquots were stored at - 80°C with glycerol (15% v/v). In parallel, continuous SEM1b cultures were maintained via regular transfers into fresh media (each recultivation incubated for ~2-3 days).

### Metagenomic analysis

Two different samples (D1B and D2B) were taken from a continuous SEM1b culture and were used for shotgun metagenomic analysis. D2B was 15 recultivations older than D1B and was used to leverage improvements in metagenome assembly and binning. From 6ml of culture, cells were pelleted by centrifugation at 14000 × *g* for 5 minutes and were kept frozen at −20°C until processing. Non-invasive DNA extraction methods were used to extract high molecular weight DNA as previously described (Kunath et al 2017). The DNA was quantified using a Qubit™ fluorimeter and the Quant-iT™ dsDNA BR Assay Kit (Invitrogen, USA) and the quality was assessed with a NanoDrop 2000 (Thermo Fisher Scientific, USA).

16S rRNA gene analysis was performed on both D1B and D2B samples. The V3-V4 hyper-variable regions of bacterial and archaeal 16S rRNA genes were amplified using the 341F/805R primer set: 5’-CCTACGGGNBGCASCAG-3’ / 5’-GACTACNVGGGTATCTAATCC-3’ (Takahashi et al 2014). The PCR was performed as previously described (Zamanzadeh et al 2016) and the sequencing library was prepared using Nextera XT Index kit according to Illumina's instructions for the MiSeq system (Illumina Inc.). MiSeq sequencing (2×300bp with paired-ends) was conducted using the MiSeq Reagent Kit v3. The reads were quality filtered (Phred ≥ Q20) and USEARCH61 (Edgar 2010) was used for detection and removal of chimeric sequences. Resulting sequences were clustered at 97% similarity into operational taxonomic units (OTUs) and taxonomically annotated with the pick_closed_reference_otus.py script from the QIIME v1.8.0 toolkit (Caporaso et al 2010) using the Greengenes database (gg_13_8). The resulting OTU table was corrected based on the predicted number of *rrs* operons for each taxon (Stoddard et al 2015).

D1B and D2B were also subjected to metagenomic shotgun sequencing using the Illumina HiSeq3000 platform (Illumina Inc) at the Norwegian Sequencing Center (NSC, Oslo, Norway). Samples were prepared with the TrueSeq DNA PCR-free preparation, and sequenced with paired-ends (2×125bp) on four lanes (two lanes per sample). Quality trimming of the raw reads was performed using cutadapt (Martin 2011), removing all bases on the 3’ end with a Phred score lower than 20 (if any present) and excluding all reads shorter than 100nt, followed by a quality filtering using the FASTX-Toolkit (http://hannonlab.cshl.edu/fastxtoolkit/). Reads with a minimum Phred score of 30 over 90% of the read length were retained. In addition, genomes from two isolated *C. proteolyticus* strains (see below) were used to decrease the data complexity and to improve the metagenomic assembly and binning. The quality-filtered metagenomic reads were mapped against the assembled strains using the BWA-MEM algorithm requiring 100% identity (Li 2013). Reads that mapped the strains were removed from the metagenomic data and the remaining reads were co-assembled using MetaSpades v3.10.0 (Nurk et al 2017) with default parameters and k-mer sizes of 21, 33, 55 and 77. The subsequent contigs were binned with Metabat v0.26.3 (Kang et al 2015) in “very sensitive mode”, using the coverage information from D1B and D2B. The quality (completeness, contamination and strain heterogeneity) of the bins (hereafter referred to as MAGs) was assessed by CheckM v1.0.7 (Parks et al 2015) with default parameters.

### Isolation of *C. proteolyticus* strains

Strains were isolated using the Hungate method (Hungate 1969). In brief: Hungate tubes were anaerobically prepared with the DSMZ medium 481 with and without agar (15g/L). Directly after being autoclaved, Hungate tubes containing agar were cooled down to 65°C and sodium sulfide nonahydrate was added. From the SEM1b culture used for D1B, 100μl were transferred to a new tube and mixed. From this new tube, 100μl was directly transferred to 10ml of fresh medium, mixed and transferred again (six transfers in total). Tubes were then cooled to 60°C for the agar to solidify, and then kept at the same temperature. After growth, single colonies were picked and transferred to liquid medium.

DNA was extracted using the aforementioned method for metagenomic DNA, with one amendment: extracted DNA was subsequently purified with DNeasy PowerClean Pro Cleanup Kit (Qiagen, USA) following manufacturer's instructions. To insure the purity of the *C. proteolyticus* colonies, visual confirmation was performed using light microscopy and long 16S rRNA genes were amplified using the primers pair 27F/1492R (Schumann 1991): 5’- AGAGTTTGATCMTGGCTCAG-3’ / 5’-TACGGYTACCTTGTTACGACTT-3’ and sequenced using Sanger technology. The PCR consisted of an initial denaturation step at 94°C for 5 min and 30 cycles of denaturation at 94°C for 1 min, annealing at 55°C for 1 min, and extension at 72°C for 1 min, and a final elongation at 72°C for 10 min. PCR products were purified using the NucleoSpin Gel and PCR Clean-up kit (Macherey-Nagel, Germany) and sent to GATC Biotech for Sanger sequencing.

The genomes of two isolated *C. proteolyticus* strains (hereafter referred to as *BWF2A* and *SW3C)* were sequenced at the Norwegian Sequencing Center (NSC, Oslo, Norway). Samples were prepared with the TrueSeq DNA PCR-free preparation and sequenced using paired-ends (2×300bp) on a MiSeq system (Illumina Inc). Quality trimming, filtering and assembly were performed as described in the aforementioned metagenomic assembly section. The raw reads were additionally mapped on assembled contigs using bowtie2 (-very-sensitive - X 1000 -I 350) and the coverage was retrieved for every nucleotide with samtool depth -a. All the contigs with an average coverage higher than 100 were selected and individually inspected for coverage discontinuity. All the contigs selected with the average coverage criterion (BWF2A: 11, SW3C: 13) look continuous in coverage and together with the Metagenome Assembled Genomes (MAGs), they were submitted to the Integrated Microbial Genomes and Microbiomes (IMG/M) system (Chen et al 2017) for genomic feature prediction and annotation (pipeline version 4.15.1). Resulting annotated open reading frames (ORFs) were retrieved, further annotated for carbohydrate-active enzymes (CAZymes) using the CAZy annotation pipeline (Lombard et al 2014), and subsequently used as a reference database for the metatranscriptomics (with exception of glycosyltransferases). The genomes for both strains and MAGs corresponding to *C. proteolyticus* were compared to the reference genome from *C. proteolyticus* DSM 5265. Using the BRIG tool (Alikhan et al 2011) for mapping and visualization, the different genomes were mapped against their pan genome generated using Roary (Page et al 2015).

### Phylogenetic analysis

A concatenated ribosomal protein phylogeny was performed on the MAGs and the isolated strains using 16 ribosomal proteins chosen as single-copy phylogenetic marker genes (RpL2, 3, 4, 5, 6, 14, 15, 16, 18, 22 and 24, and RpS3, 8, 10, 17 and 19) (Hug et al 2016). The dataset was augmented with metagenomic sequences retrieved from our previous research on the original FREVAR reactor (Hagen et al 2017) and with sequences from reference genomes identified during the 16S rRNA analysis. Each gene set was individually aligned using MUSCLE v3.8.31 (Edgar 2004) and then manually curated to remove end gaps and ambiguously aligned terminal regions. The curated alignments were concatenated and a maximum likelihood phylogeny was obtained using MEGA7 (Kumar et al 2016) with 1000 bootstrap replicates. The radial tree was visualized using iTOL (Letunic and Bork 2016). Additionally, an average nucleotide identity (ANI) comparison was performed between each MAG and their closest relative using the ANI calculator (Rodriguez-R and Konstantinidis 2016).

### Heterologous expression and purification of the GH16 enzyme

The Coprothermobacter proteolyticus BWF2A Ga0187557_1002 gene-sequence without predicted signal peptide (Petersen et al 2011) was cloned from isolated genomic DNA using the following primers; GH16_Fwd: TTAAGAAGGAGATATACTATGCTCGGCGTGAATGTGATGAATATAAGTGA; GH16_rev: AATGGTGGTGATGATGGTGCGCCTCATTTTCAAGCTTGTATACACGGACATAATC, and cloned into the pNIC-CH plasmid in E. coli TOP10 by ligation-independent cloning (Aslanidis and de Jong 1990). The transformant's sequence was verified by sequencing before transformation into OneShot^®^ E. coli BL21 Star™ cells (Thermo Fischer Scientific, Waltham, MA, USA) for expression, where 200ml Luria-broth containing 50μg/ml kanamycin was inoculated with 2ml over-night culture and incubated at 37°C, 200 rpm. Expression was induced when the culture reached an OD_600_ of 0.6, by addition of isopropyl-β-D-1-thiogalactopyranoside (IPTG). The culture was incubated at 22°C, 200rpm for 16 hours, before harvesting by centrifugation (5,000 × *g*, 10 minutes) and storage of the pellet at −80°C. The frozen pellet was transferred to 20mL buffer A (20mM Tris-HCL pH8.0, 200mM NaCl, 5mM imidazole) containing 1X BugBuster (Merck Millipore, Berlington, MA, USA), and stirred for 20 minutes at room temperature to lyse the cells. Cell debris was removed by centrifugation (30,000 *× g*, 20 minutes), and the protein was purified by immobilized metal-ion chromatography using a 5ml HisTrap FF column (GE-Healthcare, Little Chalfont, United Kingdom) pre-equilibriated with buffer A. The protein was eluted using a linear gradient to Buffer B (Buffer A with 500mM imidazole). The purity of the eluted fractions were assessed by SDS-PAGE, and the imidazole was removed from the buffer by repeated concentration and dilution using a Vivaspin (Sartorius, Göttingen, Germany) concentrator with a 10kDa cutoff. The protein concentration was determined by measured A280 and the calculated extinction coefficient.

### Biochemical characterization of the GH16 enzyme

Assays were performed in triplicate in 96-well plates, and contained 1mg/ml substrate, 20mM BisTris, pH 5.8 (50°C), and 1μM enzyme in a volume of 100μl. The reactions were pre-heated to 50°C before addition of enzyme, and were sealed before incubation for 1 hour in a Thermomixer C incubator with heated lid (Eppendorf, Hamburg, Germany). The substrates used were: barley β-glucan, carboxymethyl-curdlan, carboxymethyl-pachyman, carob galactomannan, tamarind xyloglucan, wheat arabinoxylan, larch arabinogalactan (all from Megazyme, Bray, Co. Wicklow, Ireland), and laminarin from Laminaria digitate (Sigma-Aldrich, St. Louis, MO, USA). Reactions were stopped by addition of DNS reagent (100μl, 10g/l 3,5-dinitrosalicylic acid, 300g/L Potassium sodium tartrate, 10g/L NaOH (Miller 1959)for quantification, or NaOH to a final concentration of 0.1M for product analysis. Reducing ends were quantified against a standard curve of glucose, where reactions with DNS-reagent were incubated at 95°C for 20 minutes before cooling on ice, and the absorbance was measured at 540nm. For product analysis, the reactions containing NaOH were further diluted 1:10 in water, before analysis by high-performance anion-exchange chromatography with pulsed amperometric detection (HPAEC-PAD), using a Dionex ICS-3000 system with a CarboPac PA1 column (Sunnyvale, CA, USA). Oligosaccharides were eluted using a multi-step gradient, going from 0.1M NaOH to 0.1M NaOH - 0.3M sodium acetate (NaOAc) over 35 minutes, to 0.1M NaOH - 1.0M NaOAc over 5 minutes, before going back to 0.1M NaOH over 1 minute, and reconditioning for 9 minutes at 0.1M NaOH.

### Temporal meta-omic analyses of SEM1b

A “meta-omic” time series analysis was conducted over the lifetime span of the SEM1b consortium (≈45hours). A collection of 27 replicate bottles containing ATCC medium 1943 with 10g/L of cellulose, were inoculated from the same SEM1b culture, and incubated at 65°C in parallel. For each sample time point, three culture-containing bottles were removed from the collection and processed in triplicate. Sampling occurred over nine time-points (at 0, 8, 13, 18, 23, 28, 33, 38 and 43 hours) during the SEM1b life-cycle, and are hereafter referred as T0, T1, T2, T3, T4, T5, T6, T7 and T8, respectively. DNA for 16S rRNA gene analysis was extracted (as above) from T1 to T8 and kept at −20°C until amplification and sequencing, and the analysis was performed using the protocol described above. Due to low cell biomass at the initial growth stages, sampling for metatranscriptomics was performed from T2 to T8. Sample aliquots (6 ml) were treated with RNAprotect Bacteria Reagent (Qiagen, USA) following the manufacturer's instructions and the treated cell pellets were kept at −80°C until RNA extraction.

In parallel, metadata measurements including cellulose degradation rate, monosaccharide production and protein concentration were performed over all the nine time points (T0-T8). For monosaccharide detection, 2 ml samples were taken in triplicates, centrifuged at 16000 × *g* for 5 minutes and the supernatants were filtered with 0.2μm sterile filters and boiled for 15 minutes before being stored at −20°C until processing. Solubilized sugars released during microbial hydrolysis were identified and quantified by high-performance anion exchange chromatography (HPAEC) with pulsed amperiometric detection (PAD). A Dionex ICS3000 system (Dionex, Sunnyvale, CA, USA) equipped with a CarboPac PA1 column (2 x 250 mm; Dionex, Sunnyvale, CA, USA), and connected to a guard of the same type (2 x 50 mm), was used. Separation of products was achieved using a flow rate of 0.25mL/min in a 30-minute isocratic run at 1mM KOH at 30°C. For quantification, peaks were compared to linear standard curves generated with known concentrations of selected monosaccharides (glucose, xylose, mannose, arabinose and galactose) in the range of 0.001-0.1g/L.

Total proteins measurements were taken to estimate SEM1b growth rate. Proteins were extracted following a previously described method (Hagen et al 2017) with a few modifications. Briefly, 30ml culture aliquots were centrifuged at 500 × *g* for 5 minutes to remove the substrate and the supernatant was centrifuged at 9000 × *g* for 15 minutes to pellet the cells. Cell lysis was performed by resuspending the cells in 1ml of lysis buffer (50 mM Tris-HCl, 0.1% (v/v) Triton X-100, 200 mM NaCl, 1 mM DTT, 2mM EDTA) and keeping them on ice for 30 minutes. Cells were disrupted in 3 × 60 second cycles using a FastPrep24 (MP Biomedicals, USA) and the debris were removed by centrifugation at 16000 × *g* for 15 minutes. Supernatants containing proteins were transferred into low bind protein tubes and the proteins were quantified using Bradford's method (Bradford 1976).

Because estimation of cellulose degradation requires analyzing the total content of a sample to be accurate, the measurements were performed on individual cultures that were prepared separately. A collection of 18 bottles (9 time points in duplicate) were prepared using the same inoculum described above, and grown in parallel with the 27-bottle collection used for the meta-omic analyses. For each time point, the entire sample was recovered, centrifuged at 5000 × *g* for 5 minutes and the supernatant was discarded. The resulting pellets were boiled under acidic conditions as previously described (Zhou et al 2014) and the dried weights, corresponding to the remaining cellulose, were measured.

mRNA extraction was performed in triplicate on time points T2 to T8, using previously described methods (Gifford et al 2011) with the following modifications in the processing of the RNA. The extraction of the mRNA included the addition of an *in vitro* transcribed RNA as an internal standard to estimate the number of transcripts in the natural sample compared with the number of transcripts sequenced. The standard was produced by the linearization of a pGem-3Z plasmid (Promega, USA) with ScaI (Roche, Germany). The linear plasmid was purified with a phenol/chloroform/isoamyl alcohol extraction and digestion of the plasmid was assessed by agarose gel electrophoresis. The DNA fragment was transcribed into a 994nt long RNA fragment with the Riboprobe *in vitro* Transcription System (Promega, USA) following the manufacturer's protocol. Residual DNA was removed using the Turbo DNA Free kit (Applied Biosystems, USA). The quantity and the size of the RNA standard was measured with a 2100 bioanalyzer instrument (Agilent).

Total RNA was extracted using enzymatic lysis and mechanical disruption of the cells and purified with the RNeasy mini kit following the manufacturer's protocol (Protocol 2, Qiagen, USA). The RNA standard (25ng) was added at the beginning of the extraction in every sample. After purification, residual DNA was removed using the Turbo DNA Free kit, and free nucleotides and small RNAs such as tRNAs were cleaned off with a lithium chloride precipitation solution according to ThermoFisher Scientific's recommendations. To reduce the amount of rRNAs, samples were treated to enrich for mRNAs using the MICROBExpress kit (Applied Biosystems, USA). Successful rRNA depletion was confirmed by analyzing both pre- and post-treated samples on a 2100 bioanalyzer instrument. Enriched mRNA was amplified with the MessageAmp II-Bacteria Kit (Applied Biosystems, USA) following manufacturer's instruction and sent for sequencing at the Norwegian Sequencing Center (NSC, Oslo, Norway). Samples were subjected to the TruSeq stranded RNA sample preparation, which included the production of a cDNA library, and sequenced with paired-end technology (2×125bp) on one lane of a HiSeq 3000 system.

RNA reads were assessed for overrepresented features (adapters/primers) using FastQC (www.bioinformatics.babraham.ac.uk/projects/fastqc/), and ends with detected features and/or a Phred score lower than 20 were trimmed using Trimmomatic v.0.36 (Bolger et al 2014). Subsequently, a quality filtering was applied with an average Phred threshold of 30 over a 10nt window and a minimum read length of 100nt. rRNA and tRNA were removed using SortMeRNA v.2.1b (Kopylova et al 2012). SortMeRNA was also used to isolate the reads originating from the pGem-3Z plasmid. These reads were mapped against the specific portion of the plasmid containing the Ampr gene using Bowtie2 (Langmead 2012) with default parameters and the number of reads per transcript was quantified. The remaining reads were pseudoaligned against the metagenomic dataset, augmented with the annotated strains, using Kallisto pseudo -pseudobam (Bray et al 2016). The resulting output was used to generate mapping files with bam2hits, which were used for expression quantification with mmseq (Turro et al 2011). Of the 40046 ORFs identified from the assembled SEM1b metagenome and two *C. proteolyticus* strains, 17598 (44%) were not found to be expressed, whereas 21480 (54%) were expressed and could be reliably quantified due to unique hits (reads mapping unambiguously against one unique ORF) (**Figure S1A**). The remaining 968 ORFs (2%) were expressed, but identified only with shared hits (reads mapping ambiguously against more than one ORF, resulting in an unreliable quantification of the expression of each ORF) (**Figure S1B**). Since having unique hits improves the expression estimation accuracy, the ORFs were grouped using mmseq in order to improve the precision of expression estimates, with only a small reduction in biological resolution (Turro et al 2014). The process first collapses ORFs into homologous groups if they have 100% sequence identity and then further collapses ORFs (or expression groups) if they acquire unique hits as a group (**Figure S1C**). This process generated 39146 expression groups of which 38428 (98%) were singletons (groups composed of single ORF) and 718 (2%) were groups containing more than one homologous ORF. From the initial 968 low-information ORFs, 661 (68%) became part of an expression group containing unique hits, 77 (8%) became part of ambiguous group (no unique hits) and 230 (24%) remained singletons (without unique hits). All expression groups without unique hits were then excluded from the subsequent analysis. A total of 21480 singletons and 605 multiple homologous expression groups were reliably quantified between *BWF2A, SW3C* and the SEM1b metatranscriptome (**Figure S1C**).

In order to normalize the expression estimates, sample sizes were calculated using added internal standards, as described previously (Gifford et al 2011). The number of reads mapping on the defined region of the internal standard molecule were calculated to be 4.8×10^3 +/-4.1×10^3 reads per sample out of 6.2×10^9 molecules added. Using this information, the estimated number of transcript molecules per sample was computed to be 5.1×10^12 +/- 3.7×10^12 transcripts. The resulting estimates for the sample sizes were used to scale the expression estimates from mmseq collapse and to obtain absolute expression values. During initial screening the sample T7C (time point T7, replicate C) was identified as an outlier using principle component analysis (PCA) and removed from downstream analysis.

The expression groups were clustered using hierarchical clustering with Euclidean distance. Clusters were identified using the Dynamic Tree Cut algorithm (Langfelder et al 2008) with hybrid mode, deepsplit=1 and minClusterSize=7. Eigengenes were computed for the clusters and clusters with a Pearson Correlation Coefficient (PCC) greater than 0.9 were merged. The MAG/strain enrichment of the clusters was assessed using the BiasedUrn R package. The p-values were corrected with the Benjamini-Hochberg procedure and the significance threshold was set to 0.05. Expression groups composed of multiple MAGs/strains were included in several enrichment tests.

## RESULTS AND DISCUSSION

### The SEM1b consortium is a simplistic community, co-dominated by Clostridium thermocellum and heterogeneic C. proteolyticus strains

Molecular analysis of a reproducible, cellulose-degrading and biogas-producing consortium (SEM1b) revealed a stable and simplistic population structure that contained approximately seven populations, several of which consisted of multiple strains (**Figure 2**, **Table S2-S3**). 16S rRNA gene analysis showed that the SEM1b consortium was co-dominated by OTUs affiliated to the genera *Clostridium* (52%) and *Coprothermobacter* (41%), with closest representatives identified as *Clostridium (Ruminiclostridium) thermocellum*, an uncharacterized *Clostridium spp*. and three *Coprothermobacter* phylotypes (**Table S2**). Previous meta-omic analysis on the parent Frevar reactor, revealed a multitude of numerically dominant *C. proteolyticus* strains, which created significant assembly and binning related issues (Hagen et al 2017). In this study, multiple oligotypes of *C. proteolyticus* were also found (**Table S2**). We therefore sought to isolate and recover axenic representatives to complement our meta-omic approaches, and using traditional anaerobic isolation techniques, we were successful in recovering two novel axenic strains (hereafter referred to as *BWF2A* and *SW3C)*. The genomes of *BWF2A* and *SW3C* were sequenced and assembled and subsequently incorporated into our metagenomic and metatranscriptomic analysis below.

**Figure 2.**
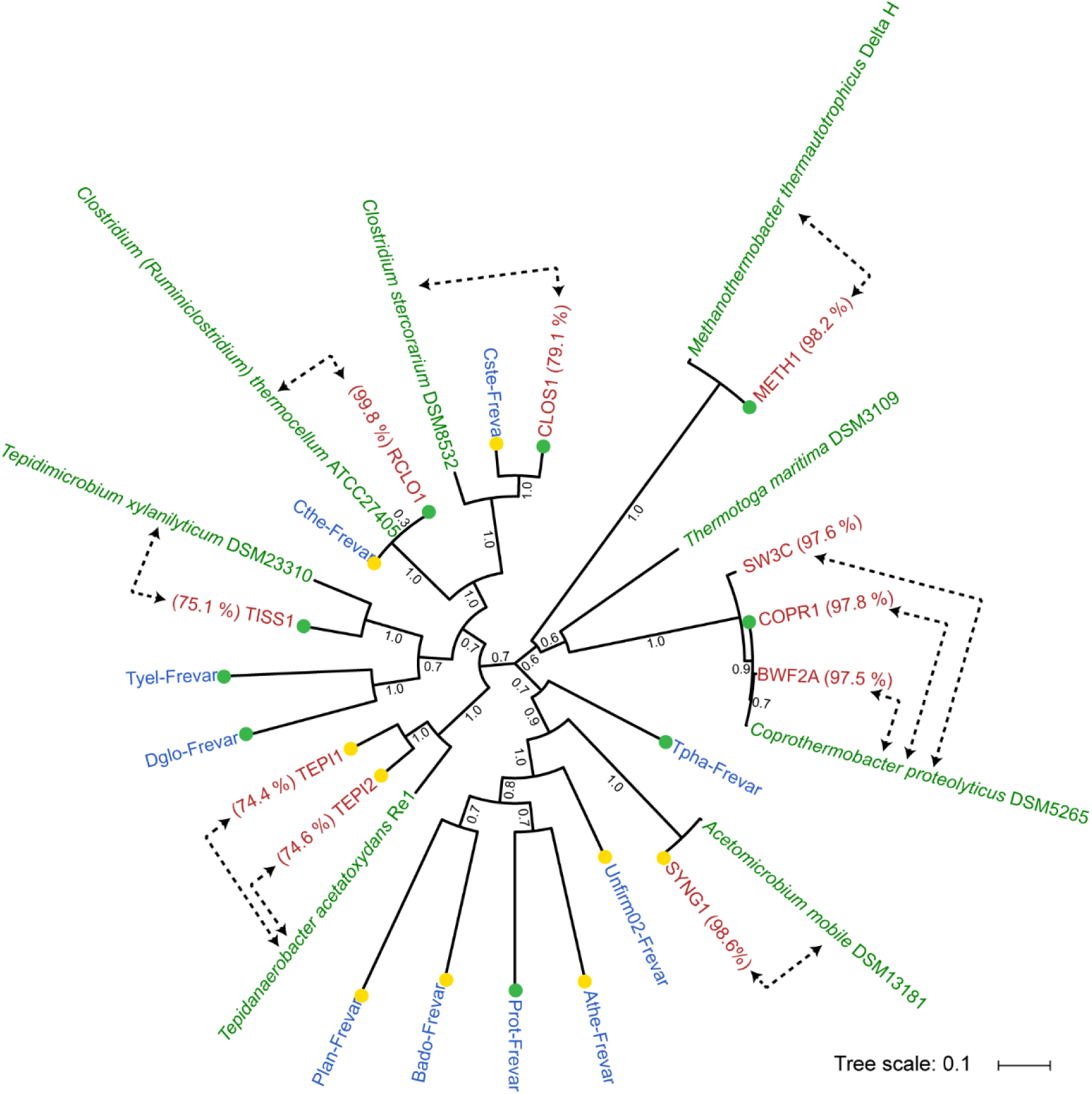
Phylogeny of *C. proteolyticus* strains and other MAGs recovered from the SEM1b consortium. Concatenated ribosomal protein tree of reference isolate genomes (green), MAGs from the previous Frevar study (blue, Hagen et al., 2017) and MAGs and isolate genomes recovered in this study (red). Average nucleotide identities (percentage indicated in parenthesis) were generated between SEM1b MAGs and their closest relative (indicated by dotted arrows). Bootstrap values are based on 1000 bootstrap replicates and the completeness of the MAGs are indicated by green (>90 %) and yellow (>80 %) colored dots.

Shotgun metagenome sequencing of two SEM1b samples (D1B and D2B), generated 290Gb (502M paired-end reads) and 264Gb (457M paired-end reads) of data, respectively. Co-assembly of both datasets using strain-depleted reads with Metaspades produced 20760 contigs totalizing 27Mbp with a maximum contig length of 603Kbp. Taxonomic binning revealed 11 MAGs and a community structure similar to the one observed by 16S analysis (**Figure 2**, **Table S3**). A total of eight MAGs exhibited high completeness (> 80%) and a low level of contamination (< 10%). Three MAGs, COPR2, COPR3 and SYNG2 corresponded to small and incomplete MAGs, although Blastp analysis suggest COPR2 and COPR3 likely represent *Coprothermobacter-affiliated* strain elements.

All near-complete MAGs (> 80%) as well as *BWF2A* and *SW3C* were phylogenetically compared against their closest relatives using average nucleotide identities (ANI) and a phylogenomic tree was constructed via analysis of 16 concatenated ribosomal proteins (**Figure 2**). One MAG was observed to cluster together with *C. proteolyticus* DSM 5265 and the two strains *BWF2A* and *SW3C* and was defined as COPR1. Two MAGs (RCLO1-CLOS1) clustered together within the *Clostridium;* RCLO1 with the well-known *C. thermocellum*, whereas CLOS1 grouped together with another *Clostridium* MAG generated from the FREVAR dataset and the isolate *C. stercorarium* (ANI: 79.1%). Both RCLO1 and CLOS1 encoded broad plant polysaccharide degrading capabilities, containing 297 and 139 carbohydrate-active enzymes (CAZymes), respectively (**Table S4**). RCLO1 in particular encoded cellulolytic (e.g. glycosyl hydrolase (GH) families GH5, GH9, GH48) and cellulosomal features (dockerins and cohesins), whereas CLOS1 appears more specialized towards hemicellulose degradation (e.g. GH3, GH10, GH26, GH43, GH51, GH130). Surprisingly, several CAZymes were also identified in COPR1 (n=65) and both *BWF2A* (n=37) and *SW3C* (n=34) at levels higher than what has previously been observed in *C. proteolyticus* DSM 5265 (n=29) (**Table S4**). Several MAGs were also affiliated with other known lineages associated with biogas processes, including *Tepidanaerobacter* (TEPI1-2), *Synergistales* (SYNG1-2), *Tissierellales* (TISS1) and Methanothermobacter (METH1).

### Novel strains of C. proteolyticus reveal acquisition of carbohydrate-active enzymes

Genome annotation of COPR1, *BWF2A* and *SW3C* identified both insertions and deletions in comparison to the only available reference genome, sequenced from the type strain DSM 5265 (**Figure 3**). Functional annotation showed that most of the genomic differences were sporadic and are predicted not to affect the metabolism of the strains. However, several notable differences were observed, which might represent a significant change in the lifestyle of the isolates. Both isolated strains lost the genes encoding flagellar proteins, although it is debatable that these genes originally conferred mobility in the type strain, as it has been previously reported as non-motile (Kersters et al 1994, Ollivier et al 1985). Interestingly, both strains acquired extra CAZymes including a particular genomic region that encoded a cluster of three CAZymes: GH16, GH3 and GH18-CBM35 (region-A, **Figure 3**). The putative function of these GHs, suggests that both *BWF2A* and *SW3C* are capable of hydrolyzing various beta-glucan linkages that are found in different hemicellulosic substrates (GH16: endo-β-1,3-1,4-glucanase; GH3: β-glucosidase). Regarding the putative GH18 encoded in both strains, it could play a role in bacterial cell wall recycling (Johnson et al 2013) as an endo-β-N-acetylglucosaminidase. Indeed, *C. proteolyticus* has previously been considered to be a scavenger of dead cells, even though this feature was mainly highlighted in term of proteolytic activities (Lü et al 2014).

**Figure 3.**
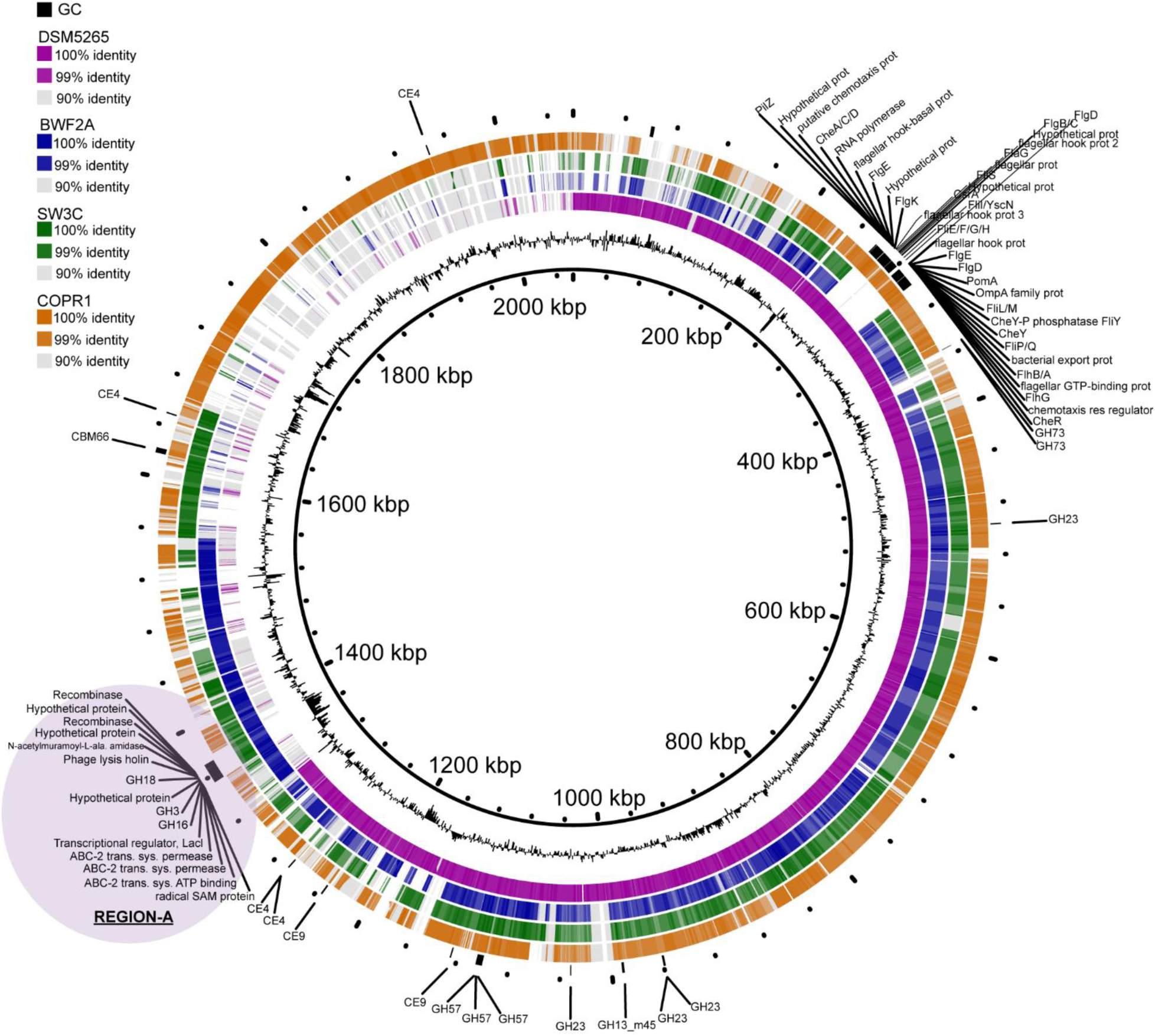
Comparative genome content of *C. proteolyticus* representatives including isolated strains, a recovered MAG (COPR1) and the reference strain DSM 5265. The innermost ring corresponds to the pangenome of the three *C. proteolyticus* spp. genomes and one MAG as produced by Roary (Page et al., 2015) and the second innermost ring represents the GC content. Outer rings represent the reference strain DSM 5265 (purple), the isolated strains BWF2A (blue) and SW3C (green) and the recovered COPR1 MAG (orange). Genes coding for carbohydrate-active enzymes (CAZymes) and flagellar proteins are indicted in black on the outermost ring. Genomic region-A is indicated by purple shading.

Taking a closer look, the region-A of CAZymes (GH16, GH3, GH18-CBM35) in *BWF2A* and *SW3C* was located on the same chromosomal cassette but organized onto two different operons with opposite directions (**Figure 4**). Comparison of the genes and their organization, revealed a high percentage of gene similarity and synteny with genome representatives from both phyla Firmicutes *(Thermoanaerobacter, Clostridium cellulolyticum* and *C. thermocellum)* and Thermotogae *(Thermosipho africanus, Fervidobacterium nodosum* and *F. gondwanense)*. Both *C. thermocellum* and *Fervidobacterium* populations were previously identified in the original Frevar reactor (Hagen et al 2017). Moreover, a truncated contig from the Frevar metagenome (Scaffold Id:Ga0101770_1036339) exhibited 99.9 % nucleotide identity to the *BWF2A* and *SW3C* genomes spanning 4.7 Kb across the CAZymes and genomic sections from both phyla (**Figure 4**), suggesting the acquirement of region-A preceded the SEM1b enrichment.

**Figure 4.**
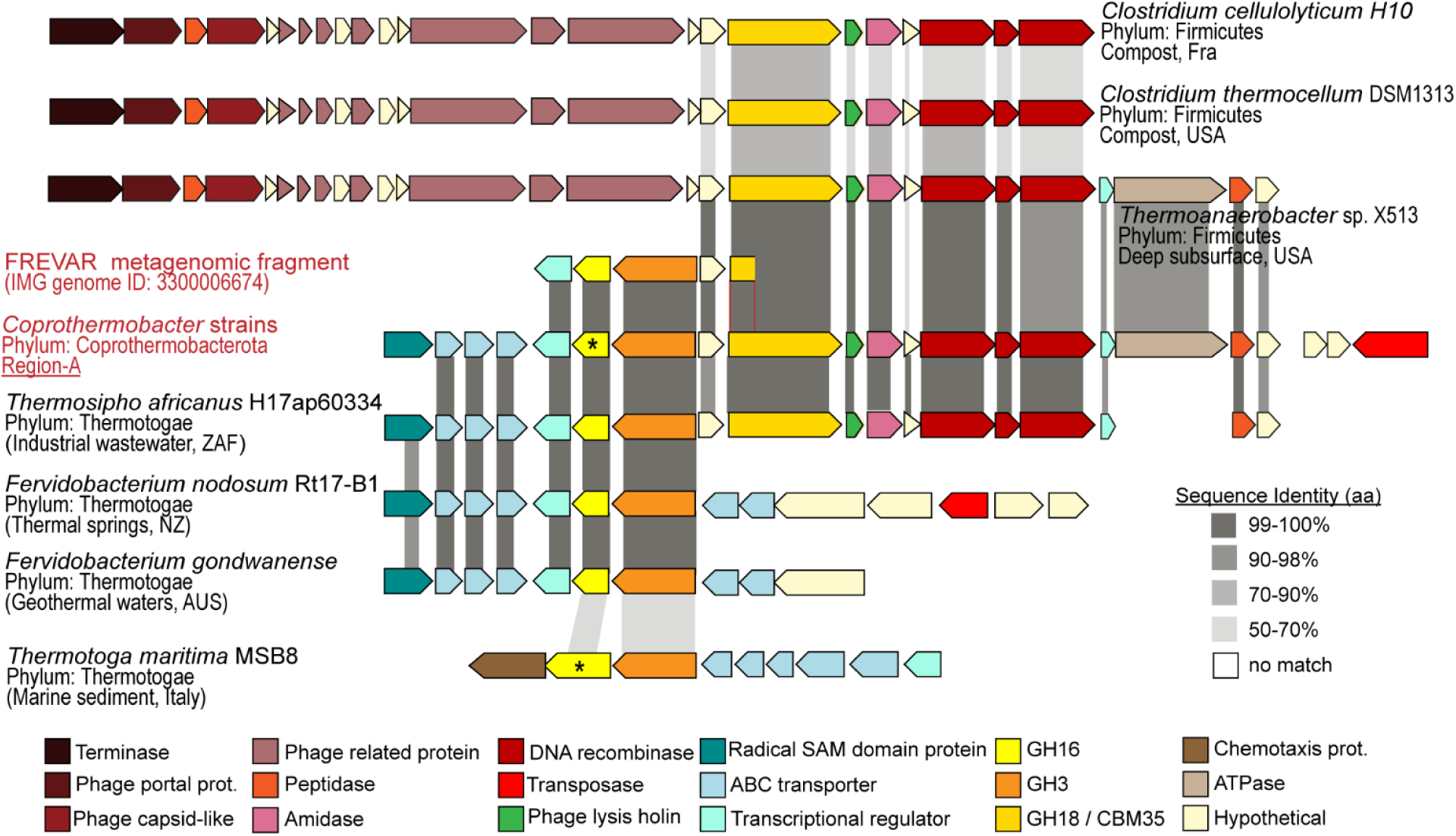
Gene synteny of CAZymes within region-A encoded in *BWF2A* and *SW3C* genomes. The gene organization of CAZymes within region-A encoded in *BWF2A* and *SW3C* (see **Figure 3**), as well as highly similar operons found in the original Frevar metagenome and isolated representatives from both phyla Firmicutes *(Thermoanaerobacter, Clostridium cellulolyticum, C. thermocellum)* and Thermotogae *(Thermosipho africanus, Fervidobacterium nodosum*, *F. gondwanense* and *Thermotoga maritima*). Grey shading between individual ORFs indicates amino acid sequence identity calculated between each query ORF (Frevar metagenome and isolates) and the reference ORF encoded in region-A from *BWF2A* and *SW3C* (identical in both strains). Asterisk denotes biochemically characterized GH16 enzymes, including the *C. proteolyticus* representative from this study and a laminarinase from *Thermotoa maritima* MSB8 that has previously been reported (Jeng et al 2011).

Examination of the flanking regions surrounding the CAZymes in region-A, reveals the presence of an incomplete prophage composed of a phage lysis holin and two recombinases located downstream (**Figure 3**, **Figure 4**). Further comparisons revealed that only the Firmicutes-lineages encoded the same prophage together with an additional terminase, phage-capsid like proteins and more phage-related components on the 5’ region (**Figure 4**). Because of the high sequence homology and the presence of phage-genes in the surrounding, we hypothesized that the origin of region-A in *BWF2A* and *SW3C*, is the result of phage-mediated HGT. Most likely, the operon from Firmicutes-affilaited lineages (e.g. *Thermoanaerobacter* and *C. thermocellum)* was transferred first due to the presence of its complete phage and generated a hot spot for further HGT for the GH16-GH3 encoding operon originating from Thermotogae-affiliated lineages (**Figure 4**). Interestingly, *T. africanus* also encoded a syntenous region that covered Region-A in both *BWF2A* and *SW3C* almost in its entirety (**Figure 4**), creating an alternative possibility that vertical gene transfer may also have played a role towards the evolution of this operon in *Coprothermobacter*. Gene transfer within anaerobic digesters has been reported for antibiotic resistance genes (Miller et al 2016), whereas HGT of CAZymes have been detected previously among gut microbiota (Hehemann et al 2010, Ricard et al 2006, Song et al 2016). Since many microbes express only a specific array of carbohydrate-degrading capabilities, bacteria that acquire CAZymes from gene transfer events may gain additional capacities and consequently, a selective growth advantage (Modi et al 2013).

In response to our discovery of *C. proteolyticus* CAZyme acquisition, we attempted to cultivate our axenic strains in minimal media containing only hemicellulosic substrates (pachyman, curdlan, barley beta-glucan) as a sole carbon source. However, no growth was observed for either *BWF2A* or *SW3C* in polysaccharide-supplemented media that was without yeast extract. These results were consistent with the few available studies on type strain DSM 5265, which have shown weak and slow growth on proteins and monomeric sugars, and only in the presence of pluralistic organic compounds found in yeast extract and rumen fluid (Kersters et al 1994, Ollivier et al 1985). Growth was observed in *BWF2A/SW3C* cultures with both yeast extract and polysaccharide substrates, however we detected no increased levels of growth, indicating that in isolation our *C. proteolyticus* strains may require specific undefined cofactor(s) or collaborative microbial partners to support the activity encoded by their acquired CAZymes.

In lieu of axenic *C. proteolyticus* cultivation data to support a saccharolytic lifestyle, we biochemically interrogated the GH16 encoded in region-A (**Figure 4**). The catalytic domain was synthesized and expressed in *Escherichia coli*, followed by protein purification. As expected the GH16 demonstrated endoglucanase activity on β-1,3 (pachyman, curdlan, laminarin) and β-1,3-1,4 (Barley) substrates (**Figure S2A**), which supports our hypothesis that the CAZymes in region-A have transferred the ability of *BWF2A* or *SW3C* to degrade polysaccharides. Against all β-glucan substrates, GH16 hydrolysis generated a large fraction of glucose (**Figure S2B**), which has been shown to be readily fermented by *C. proteolyticus* (Kersters et al 1994, Ollivier et al 1985).

### C. proteolyticus expresses CAZymes and is implicit in collaborative polysaccharide degradation within the SEM1b consortium

Whilst we confirmed that the acquired *C. proteolyticus* GH16 is functionally active, we also sought to better understand the role(s) played by it and other *C. proteolyticus* CAZymes in a saccharolytic consortium, by analyzing the temporal metatranscriptome of SEM1b over a complete life cycle. 16S rRNA gene analysis of eight time points (T1-8) over a 43hr period reaffirmed that *C. thermocellum-* and *C. proteolyticus-affiliated* populations dominate SEM1b over time (**Figure 5A**). Highly similar genes from different MAGs/genomes were grouped together in order to obtain “expression groups” with discernable expression profiles (see Methods and **Figure S1A/B**). A total of 274 singleton CAZyme expression groups and 8 multiple ORF groups were collectively detected in the two *C. proteolyticus* strains and MAGs suspected of contributing to polysaccharide degradation (RCLO1, CLOS1, COPR1-3, and TISS1, **Figure S1D**, **Table S5**). In several instances, expressed CAZymes from *BWF2A* and *SW3C* could not be resolved between the two strains and/or the COPR1 MAG. For example, all GHs within region-A could be identified as expressed by at least one of the isolated strains but could not be resolved further between the strains.

**Figure 5.**
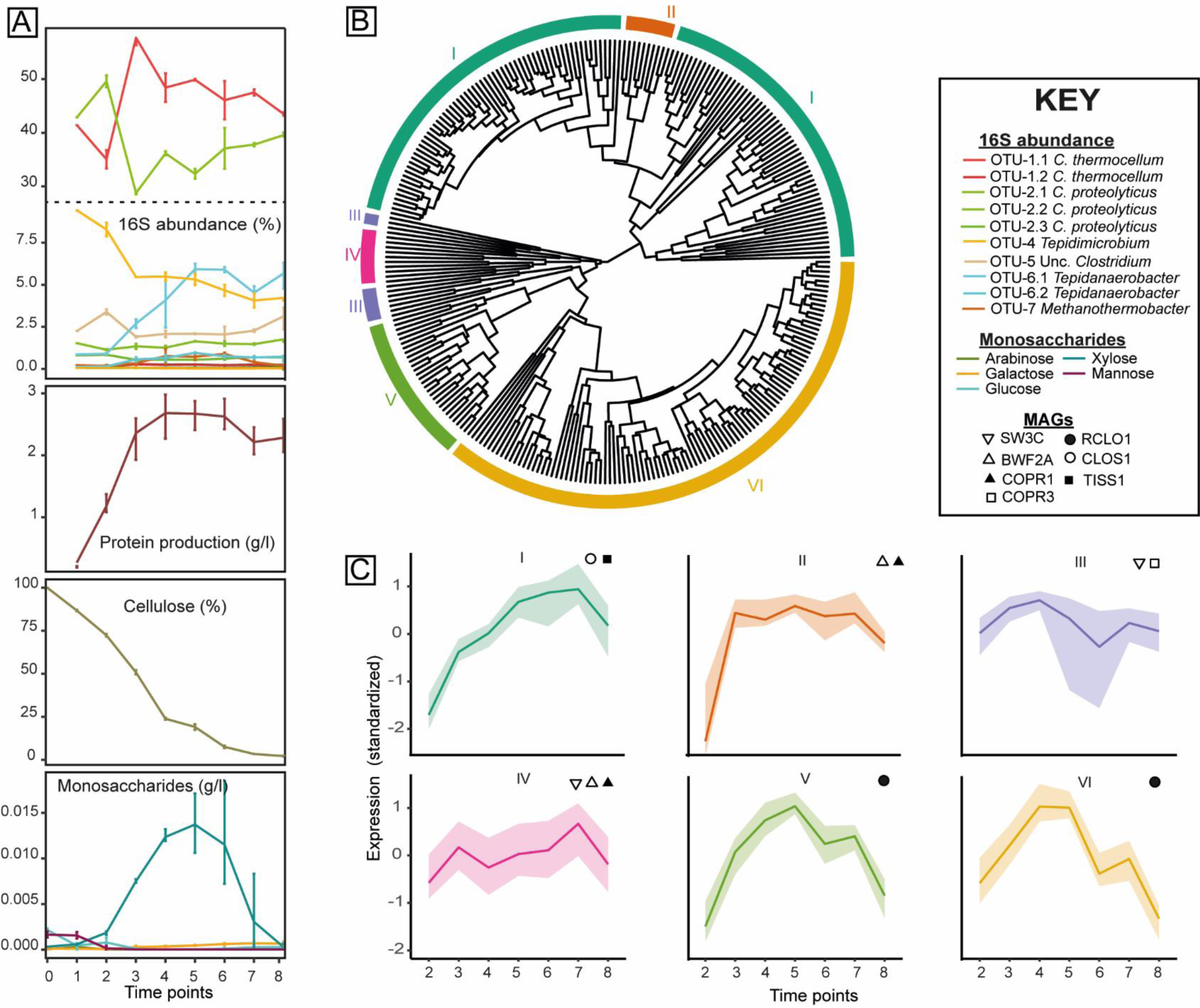
Temporal meta-analysis of the SEM1b consortium. (A) 16S rRNA gene amplicon and metadata analysis was performed over a 43-hour period, which was segmented into 9 time-points. OTU IDs are detailed in **Table S2**. Cellulose degradation rate, monosaccharide accumulation and growth rate (estimated by total protein concentration) is presented. (B) Gene expression dendrogram and clustering of CAZymes from BWF2A, SW3C and MAGs: RCLO1, CLOS1, COPR1-3, and TISS1. Six expression clusters (I-VI) are displayed in different colors on the outer ring. (C) Clusters I-VI show characteristic behaviors over time summarized by the median (solid line) and the shaded area between the first and third quartile of the standardized expression. Bacteria that are statistically enriched (p-value < 0.05) in the clusters are displayed in the subpanels.

From the CAZymes subset of expression groups, a cluster analysis was performed to reveal six expression clusters (I-VI, **Figure 5B**). Clusters II, III and IV were enriched with *C. proteolyticus-affiliated* MAGs and isolated strains. Clusters III and IV comprised of 10 and 11 expression groups, respectively, and followed a similar profile over time (**Figure 5C**), increasing at earlier stages (T2-3) and again at later stationary/death stages (T6-8). Cluster II (10 expression groups) was slightly variant and increased more rapidly at T2 and sustained high levels over the course of SEM1b. All three clusters consisted of CAZymes targeting linkages associated with N-acetylglucosamine (CE9), and peptidoglycan (CE4, GH23, GH73), suggesting a role in bacterial cell wall hydrolysis (**Table S5**). This hypothesis was supported by 16S rRNA gene data, which illustrated that *C. proteolyticus-*affiliated populations (OTU2) were high at initial stages of the SEM1b life-cycle when cell debris was likely present in the inoculum that was sourced from the preceding culture at stationary phase (**Figure 5A**). At T2, the abundance of *C. thermocellum-affiliated* populations (OTU-1) was observed to outrank *C. proteolyticus* as the community predictably shifted to cellulose-utilization. However, towards stationary phase (T6-8) when dead cell debris is expected to be increasing, expression levels in clusters II, III and IV were maintained at high levels (**Figure 5B**), which was consistent with high *C. proteolyticus* 16S rRNA gene abundance at the same time-points.

Clusters V and VI comprised 28 and 101 expression groups (respectively), and were enriched with the RCLO1 MAG that was closely related to *C. thermocellum*. As expected, numerous expressed genes in cluster V and VI were inferred in cellulosome assembly (via dockerin domains) as well as cellulose (e.g. GH5, GH9, GH44, GH48, CBM3) and hemicellulose (e.g. GH10, GH11, GH26, GH43, GH74) hydrolysis (**Table S5**). Both clusters increased throughout the consortium's exponential phase (time points T1-4, **Figure 5A**), whilst 16S rRNA data also shows *C. thermocellum-affiliated* populations at high levels during the same stages (**Figure 5A**).

Cluster I was determined as the largest with 121 expression groups, and was particularity enriched with CLOS1, which expressed many genes involved in hemicellulose deconstruction (e.g. GH3, GH10, GH29, GH31, GH43 and GH130) and carbohydrate deacetylation (e.g. CE4, CE7, CE8, CE9, CE12, CE15) (**Table S5**). CAZymes from both *BWF2A* and *SW3C* were also expressed in cluster I including the functionally active GH16 and GH3-encoding ORFs from region-A, which reaffirms our earlier predictions that certain *C. proteolyticus* populations in SEM1b are capable of degrading hemicellulosic substrates. The expression profile of cluster I over time was observed to slightly lag after cluster V and VI (**Figure 5**), suggesting that hemicellulases in cluster I genes are expressed once the hydrolytic effects of the RCLO1-cellulosome (expressed in cluster V and VI) have liberated hemicellulosic substrates (Zverlov et al 2005b). Although *C. thermocellum* cannot readily utilize other carbohydrates besides glucose and longer glucans (Demain et al 2005), the cellulosome is composed of a number of hemicellulolytic enzymes such as GH10 and GH11 endoxylanases, GH26 mannanases, GH74 xyloglucanases and GH43 arabinanases/xylosidases (Zverlov et al 2005a), which are involved in the deconstruction of the underlying cellulose-hemicellulose matrix (Zverlov et al 2005b). Interestingly, RCLO1 representatives of GH10, GH11, GH5, GH9, GH16 and GH43 were all expressed in the additional RCLO1-enriched cluster V and are presumably acting on the hemicellulose fraction present in the spruce-derived cellulose (Chylenski et al 2017). Furthermore, detection of hydrolysis products (**Figure 5A**), revealed that xylose increased significantly at T5-7, indicating that hemicellulosic polymers containing beta-1-4-xylan were likely available at these stages. Cluster V exhibited a similar profile to the other RCLO1-enriched cluster (Cluster VI), however its high expression levels were extended to T7, consistent with our observed levels of xylose release (**Figure 5C**)

An additional GH16 from RCLO1 was also expressed in SEM1b cluster V, which has 99.5 % amino acid sequence identity to Lic16A, a biochemically characterized endoglucanase that exerts specific β-1,3 activity similar to the *BWF2A/SW3C* GH16 that we report here. Notably, Lic16A is a cell wall anchored, non-cellulosomal CAZyme that is believed to enable *C. thermocellum* to grow exclusively on β-1,3-glucans (Fuchs et al 2003). All in all, the SEM1b expression data shows sequential community progression that co-ordinates putative hydrolysis of cellulose and hemicellulosic substrates as well as carbohydrates that are found in the microbial cell wall. In particular, *C. proteolyticus* populations in SEM1b were suspected to play key roles degrading microbial cell wall carbohydrates as well as hemicellulosic substrates, possibly in cooperation or in parallel to other clostridium populations at the later stages of the SEM1b growth cycle.

## CONCLUSIONS

Unraveling the interactions occurring in a complex microbial community composed of closely related species or strains is an arduous task. Here, we have leveraged culturing techniques, metagenomics, time-resolved metatranscriptomics and enzymology to describe a novel *C. proteolyticus* population that is comprised of closely related strains that have acquired CAZymes via HGT and putatively evolved to incorporate a saccharolytic lifestyle. The co-expression patterns of *C. proteolyticus* CAZymes in clusters II, III and IV supports the adaptable role of this bacterium as a scavenger that is able to hydrolyze cell wall polysaccharides during initial phases of growth and in the stationary / death phase, when available sugars are low. Moreover, the acquisition of biochemically-verified hemicellulases by *C. proteolyticus*, and their co-expression in cluster I at time points when hemicellulose is available, further enhances its metabolic versatility and provides substantial evidence as to why this population dominates thermophilic reactors on a global scale, even when substrates are poor in protein.

## DATA AVAILABILITY

All sequencing reads have been deposited in the sequence read archive (SRP134228), with specific numbers listed in **Table S6**. All microbial genomes are publicly available on JGI under the analysis project numbers listed in **Table S6**.

## ACKNOWLEDGEMENTS

We are grateful for support from The Research Council of Norway (FRIPRO program, PBP: 250479 / NorZymeD, VGHE: 221568), as well as the European Research Commission Starting Grant Fellowship (awarded to PBP; 336355 - MicroDE). The sequencing service was provided by the Norwegian Sequencing Centre (www.sequencing.uio.no), a national technology platform hosted by the University of Oslo and supported by the “Functional Genomics” and “Infrastructure” programs of the Research Council of Norway and the Southeastern Regional Health Authorities.

## REFERENCES

Alexiev A, Coil DA, Badger JH, Enticknap J, Ward N, Robb FT et al (2014). Complete Genome Sequence of Coprothermobacter proteolyticus DSM 5265. Genome Announcements 2.

Alikhan NF, Petty NK, Ben Zakour NL, Beatson SA (2011). BLAST Ring Image Generator (BRIG): Simple prokaryote genome comparisons. BMC Genomics 12.

Aslanidis C, de Jong PJ (1990). Ligation-independent cloning of PCR products (LIC-PCR). Nucleic Acids Research 18: 6069–6074.

Bendall ML, Stevens SLR, Chan LK, Malfatti S, Schwientek P, Tremblay J et al (2016). Genome-wide selective sweeps and gene-specific sweeps in natural bacterial populations. ISME Journal 10: 1589–1601.

Biller SJ, Berube PM, Lindell D, Chisholm SW (2015). Prochlorococcus: The structure and function of collective diversity. Nature Reviews Microbiology 13: 13–27.

Bolger AM, Lohse M, Usadel B (2014). Trimmomatic: A flexible trimmer for Illumina sequence data. Bioinformatics 30: 2114–2120.

Bradford MM (1976). A rapid and sensitive method for the quantitation of microgram quantities of protein utilizing the principle of protein-dye binding. Analytical Biochemistry 72: 248–254.

Bray NL, Pimentel H, Melsted P, Pachter L (2016). Near-optimal probabilistic RNA-seq quantification. Nature Biotechnology 34: 525–527.

Bron PA, Van Baarlen P, Kleerebezem M (2012). Emerging molecular insights into the interaction between probiotics and the host intestinal mucosa. Nature Reviews Microbiology 10: 66–78.

Caporaso JG, Kuczynski J, Stombaugh J, Bittinger K, Bushman FD, Costello EK et al (2010). QIIME allows analysis of high-throughput community sequencing data. Nature methods 7: 335–336.

Chen IMA, Markowitz VM, Chu K, Palaniappan K, Szeto E, Pillay M et al (2017). IMG/M: Integrated genome and metagenome comparative data analysis system. Nucleic Acids Research 45: D507–D516.

Chylenski P, Petrovic DM, Müller G, Dahlström M, Bengtsson O, Lersch M et al (2017). Enzymatic degradation of sulfite-pulped softwoods and the role of LPMOs. Biotechnology for Biofuels 10: 1–13.

Demain AL, Newcomb M, Wu JHD, Demain AL, Newcomb M, Wu JHD (2005). Cellulase, Clostridia, and Ethanol. Microbiology and Molecular Biology Reviews 69: 124–154.

Edgar RC (2004). MUSCLE: Multiple sequence alignment with high accuracy and high throughput. Nucleic Acids Research 32: 1792–1797.

Edgar RC (2010). Search and clustering orders of magnitude faster than BLAST. Bioinformatics 26: 2460–2461.

Ellegaard KM, Engel P (2016). Beyond 16S rRNA community profiling: Intra-species diversity in the gut microbiota. Frontiers in Microbiology 7: 1–16.

Etchebehere C, Pavan ME, Zorzópulos J, Soubes M, Muxí L (1998). Coprothermobacter platensis sp. nov., a new anaerobic proteolytic thermophilic bacterium isolated from an anaerobic mesophilic sludge. International journal of systematic bacteriology 48: 1297–1304.

Fuchs K-P, Zverlov VV, Velikodvorskaya GA, Lottspeich F, Schwarz WH (2003). Lic16A of Clostridium thermocellum, a non-cellulosomal, highly complex endo-β-1,3-glucanase bound to the outer cell surface. Microbiology 149: 1021–1031.

Gifford SM, Sharma S, Rinta-Kanto JM, Moran MA (2011). Quantitative analysis of a deeply sequenced marine microbial metatranscriptome. ISME Journal 5: 461–472.

Gill SR, Fouts DE, Archer GL, Mongodin EF, Deboy RT, Ravel J et al (2005). Insights on Evolution of Virulence and Resistance from the Complete Genome Analysis of an Early Methicillin-Resistant Staphylococcus aureus Strain and a Biofilm-Producing Methicillin-Resistant Staphylococcus epidermidis Strain. J Bacteriol 187: 2426–2438.

González-Torres P, Pryszcz LP, Santos F, Martínez-García M, Gabaldón T, Antón J (2015). Interactions between Closely Related Bacterial Strains Are Revealed by Deep Transcriptome Sequencing. Applied and Environmental Microbiology 81: 8445–8456.

Hagen LH, Frank JA, Zamanzadeh M, Eijsink VGH, Pope PB, Horn SJ et al (2017). Quantitative metaproteomics highlight the metabolic contributions of uncultured phylotypes in a thermophilic anaerobic digester. Applied and Environmental Microbiology 83.

Hehemann JH, Correc G, Barbeyron T, Helbert W, Czjzek M, Michel G (2010). Transfer of carbohydrate-active enzymes from marine bacteria to Japanese gut microbiota. Nature 464: 908–912.

Hug LA, Baker BJ, Anantharaman K, Brown CT, Probst AJ, Castelle CJ et al (2016). A new view of the tree of life. Nature Microbiology 1: 1–6.

Hungate RE (1969). Chapter IV A Roll Tube Method for Cultivation of Strict Anaerobes. In: Norris JR, Ribbons DWBTMiM (eds). Methods in Microbiology. Academic Press. pp 117–132.

Hunt DE, David LA, Gevers D, Preheim SP, Alm EJ, Polz MF (2008). Resource Partitioning and Sympatric Differentiation Among Closely Related Bacterioplankton. Science 320: 1081 LP-1085.

Jeng W-Y, Wang N-C, Lin C-T, Shyur L-F, Wang AHJ (2011). Crystal Structures of the Laminarinase Catalytic Domain from Thermotoga maritima MSB8 in Complex with Inhibitors: essential residues for β-1,3 and β-1,4 glucan selection. The Journal of Biological Chemistry 286: 45030–45040.

Johnson JW, Fisher JF, Mobashery S (2013). Bacterial cell wall recycling. Annals of the new york academy 1277: 54–75.

Kang DD, Froula J, Egan R, Wang Z (2015). MetaBAT, an efficient tool for accurately reconstructing single genomes from complex microbial communities. PeerJ 3: e1165–e1165.

Kashtan N, Roggensack SE, Rodrigue S, Thompson JW, Biller SJ, Coe A et al (2014). Single-Cell Genomics Reveals Hundreds of coexisting subpopulations in wild Prochlorococcus. Science (New York, NY) 344: 416–420.

Kersters I, Maestrojuan GM, Torck U, Vancanneyt M, Kersters K, Verstraete W (1994). Isolation of *Coprothermobacter proteolyticus* from an Anaerobic Digest and Further Characterization of the Species. Syst Appl Microbiol 17: 289–295.

Kopylova E, Noé L, Touzet H (2012). SortMeRNA: Fast and accurate filtering of ribosomal RNAs in metatranscriptomic data. Bioinformatics 28: 3211–3217.

Koskella B, Vos M (2015). Adaptation in Natural Microbial Populations. Annual Review of Ecology, Evolution, and Systematics 46: 503–522.

Kumar S, Stecher G, Tamura K (2016). MEGA7: Molecular Evolutionary Genetics Analysis Version 7.0 for Bigger Datasets. Molecular biology and evolution 33: 1870–1874.

Kunath BJ, Bremges A, Weimann A, McHardy AC, Pope PB (2017). Metagenomics and CAZyme Discovery. In: Abbott DW, Lammerts van Bueren A (eds). Protein-Carbohydrate Interactions: Methods and Protocols. Springer New York: New York, NY. pp 255–277.

Langfelder P, Zhang B, Horvath S (2008). Defining clusters from a hierarchical cluster tree: The Dynamic Tree Cut package for R. Bioinformatics 24: 719–720.

Langmead (2012). Fast gapped-read alignment with Bowtie 2. Nature methods 9: 357–359.

Letunic I, Bork P (2016). Interactive tree of life (iTOL) v3: an online tool for the display and annotation of phylogenetic and other trees. Nucleic acids research 44: W242–W245.

Li H (2013). Aligning sequence reads, clone sequences and assembly contigs with BWA-MEM. arxiv 00: 1–3.

Lombard V, Golaconda Ramulu H, Drula E, Coutinho PM, Henrissat B (2014). The carbohydrate-active enzymes database (CAZy) in 2013. Nucleic Acids Research 42: D490–D495.

Lü F, Bize A, Guillot A, Monnet V, Madigou C, Chapleur O et al (2014). Metaproteomics of cellulose methanisation under thermophilic conditions reveals a surprisingly high proteolytic activity. ISME Journal 8: 88–102.

Martin M (2011). Cutadapt removes adapter sequences from high-throughput sequencing reads. EMBnetjournal 17: 10–10.

McLoughlin K, Schluter J, Rakoff-Nahoum S, Smith AL, Foster KR (2016). Host Selection of Microbiota via Differential Adhesion. Cell Host and Microbe 19: 550–559.

Miller GL (1959). Use of Dinitrosalicylic Acid Reagent for Determination of Reducing Sugar. Analytical Chemistry 31: 426–428.

Miller JH, Novak JT, Knocke WR, Pruden A (2016). Survival of antibiotic resistant bacteria and horizontal gene transfer control antibiotic resistance gene content in anaerobic digesters. Frontiers in Microbiology 7: 1–11.

Modi SR, Lee HH, Spina CS, Collins JJ (2013). Antibiotic treatment expands the resistance reservoir and ecological network of the phage metagenome. Nature 499: 219–222.

Nurk S, Meleshko D, Korobeynikov A, Pevzner PA (2017). MetaSPAdes: A new versatile metagenomic assembler. Genome Research 27: 824–834.

Ochman H, Lawrence JG, Grolsman EA (2000). Lateral gene transfer and the nature of bacterial innovation. Nature 405: 299–304.

Ollivier BM, Mah Ra, Ferguson TJ, Boone DR, Garcia JL, Robinson R (1985). Emendation of the Genus Thermobacteroides: Thermobacteroides proteolyticus sp. nov., a proteolytic acetogen from a methanogenic enrichment. International Journal of Systematic Bacteriology 35: 425–428.

Page AJ, Cummins CA, Hunt M, Wong VK, Reuter S, Holden MTG et al (2015). Roary: Rapid large-scale prokaryote pan genome analysis. Bioinformatics 31: 3691–3693.

Parks DH, Imelfort M, Skennerton CT, Hugenholtz P, Tyson GW (2015). CheckM: Assessing the quality of microbial genomes recovered from isolates, single cells, and metagenomes. Genome Research 25: 1043–1055.

Petersen TN, Brunak S, von Heijne G, Nielsen H (2011). SignalP 4.0: discriminating signal peptides from transmembrane regions. Nature Methods 8: 785.

Ricard G, McEwan NR, Dutilh BE, Jouany JP, Macheboeuf D, Mitsumori M et al (2006). Horizontal gene transfer from bacteria to rumen ciliates indicates adaptation to their anaerobic, carbohydrates-rich environment. BMC Genomics 7: 1–13.

Rodriguez-R LM, Konstantinidis KT (2016). The enveomics collection: a toolbox for specialized analyses of microbial genomes and metagenomes. PeerJ Preprints 4: e1900v1901.

Rodriguez-Valera F, Martin-Cuadrado A-B, Rodriguez-Brito B, Pašić L, Thingstad TF, Rohwer F et al (2009). Explaining microbial population genomics through phage predation. Nature Reviews Microbiology 7: 828–828.

Rødsrud G, Lersch M, Sjöde A (2012). History and future of world's most advanced biorefinery in operation. Biomass and Bioenergy 46: 46–59.

Rosenzweig RF, Sharp RR, Treves DS, Adams J (1994). Microbial evolution in a simple unstructured environment: Genetic differentiation in Escherichia coli. Genetics 137: 903–917.

Schloissnig S, Arumugam M, Sunagawa S, Mitreva M, Tap J, Zhu A et al (2013). Genomic variation landscape of the human gut microbiome. Nature 493: 45–50.

Schumann P (1991). Nucleic Acid Techniques in Bacterial Systematics (Modern Microbiological Methods). Journal of Basic Microbiology 31: 479–480.

Shapiro BJ, Timberlake SC, Szabó G, Polz MF, Alm EJ (2012). Population Genomics of Early Differentiation of Bacteria. Science 336: 48–51.

Sharon I, Morowitz MJ, Thomas BC, Costello EK, Relman DA, Banfield JF (2013). Time series community genomics analysis reveals rapid shifts in bacterial species, strains, and phage during infant gut colonization. Genome Research 23: 111–120.

Siezen RJ, Tzeneva VA, Castioni A, Wels M, Phan HTK, Rademaker JLW et al (2010). Phenotypic and genomic diversity of Lactobacillus plantarum strains isolated from various environmental niches. Environmental Microbiology 12: 758–773.

Solheim M, Aakra Å, Snipen LG, Brede DA, Nes IF (2009). Comparative genomics of Enterococcus faecalis from healthy Norwegian infants. BMC Genomics 10: 1–11.

Song T, Xu H, Wei C, Jiang T, Qin S, Zhang W et al (2016). Horizontal Transfer of a Novel Soil Agarase Gene from Marine Bacteria to Soil Bacteria via Human Microbiota. Scientific Reports 6: 1–10.

Spanogiannopoulos P, Bess EN, Carmody RN, Turnbaugh PJ (2016). The microbial pharmacists within us: A metagenomic view of xenobiotic metabolism. Nature Reviews Microbiology 14: 273–287.

Stoddard SF, Smith BJ, Hein R, Roller BRK, Schmidt TM (2015). rrnDB: Improved tools for interpreting rRNA gene abundance in bacteria and archaea and a new foundation for future development. Nucleic Acids Research 43: D593–D598.

Takahashi S, Tomita J, Nishioka K, Hisada T, Nishijima M (2014). Development of a prokaryotic universal primer for simultaneous analysis of Bacteria and Archaea using next-generation sequencing. PLoS ONE 9.

Tandishabo K, Nakamura K, Umetsu K, Takamizawa K (2012). Distribution and role of Coprothermobacter spp. in anaerobic digesters. Journal of Bioscience and Bioengineering 114: 518–520.

Tettelin H, Masignani V, Cieslewicz MJ, Donati C, Medini D, Ward NL et al (2005). Genome analysis of multiple pathogenic isolates of Streptococcus agalactiae: Implications for the microbial "pan-genome". Proceedings of the National Academy of Sciences 102: 13950–13955.

Treangen TJ, Rocha EPC (2011). Horizontal transfer, not duplication, drives the expansion of protein families in prokaryotes. PLoS Genetics 7.

Truong DT, Tett A, Pasolli E, Huttenhower C, Segata N (2017). Microbial strain-level population structure & genetic diversity from metagenomes. Genome Research 27: 626–638.

Turro E, Su SY, Gonçalves Â, Coin LJM, Richardson S, Lewin A (2011). Haplotype and isoform specific expression estimation using multi-mapping RNA-seq reads. Genome Biology 12: 1–15.

Turro E, Astle WJ, Tavaré S (2014). Flexible analysis of RNA-seq data using mixed effects models. Bioinformatics 30: 180–188.

Zamanzadeh M, Hagen LH, Svensson K, Linjordet R, Horn SJ (2016). Anaerobic digestion of food waste - Effect of recirculation and temperature on performance and microbiology. Water Research 96: 246–254.

Zelezniak A, Andrejev S, Ponomarova O, Mende DR, Bork P, Patil KR (2015). Metabolic dependencies drive species co-occurrence in diverse microbial communities. Proceedings of the National Academy of Sciences 112: 6449–6454.

Zhou Y, Pope PB, Li S, Wen B, Tan F, Cheng S et al (2014). Omics-based interpretation of synergism in a soil-derived cellulose-degrading microbial community. Scientific Reports 4: 1–6.

Zunino P, Piccini C, Legnani-Fajardo C (1994). Flagellate and non-flagellate Proteus mirabilis in the development of experimental urinary tract infection. Microbial Pathogenesis 16: 379–385.

Zverlov VV, Kellermann J, Schwarz WH (2005a). Functional subgenomics of Clostridium thermocellum cellulosomal genes: Identification of the major catalytic components in the extracellular complex and detection of three new enzymes. Proteomics 5: 3646–3653.

Zverlov VV, Schantz N, Schmitt-Kopplin P, Schwarz WH (2005b). Two new major subunits in the cellusome of Clostridium thermocellum: Xyloglucanase Xgh74A and endoxylanase Xyn10D. Microbiology 151: 3395–3401.

