## Supplementary Materials for "From proteins to polysaccharides: lifestyle and genetic evolution of *Coprothermobacter proteolyticus*" (from another version of this preprint: version 2, 2018-03-17)

This PDF file includes:

I. Supplementary Figures and Legends
II. Supplementary Tables and Legends
III. References cited in the Supplementary Material

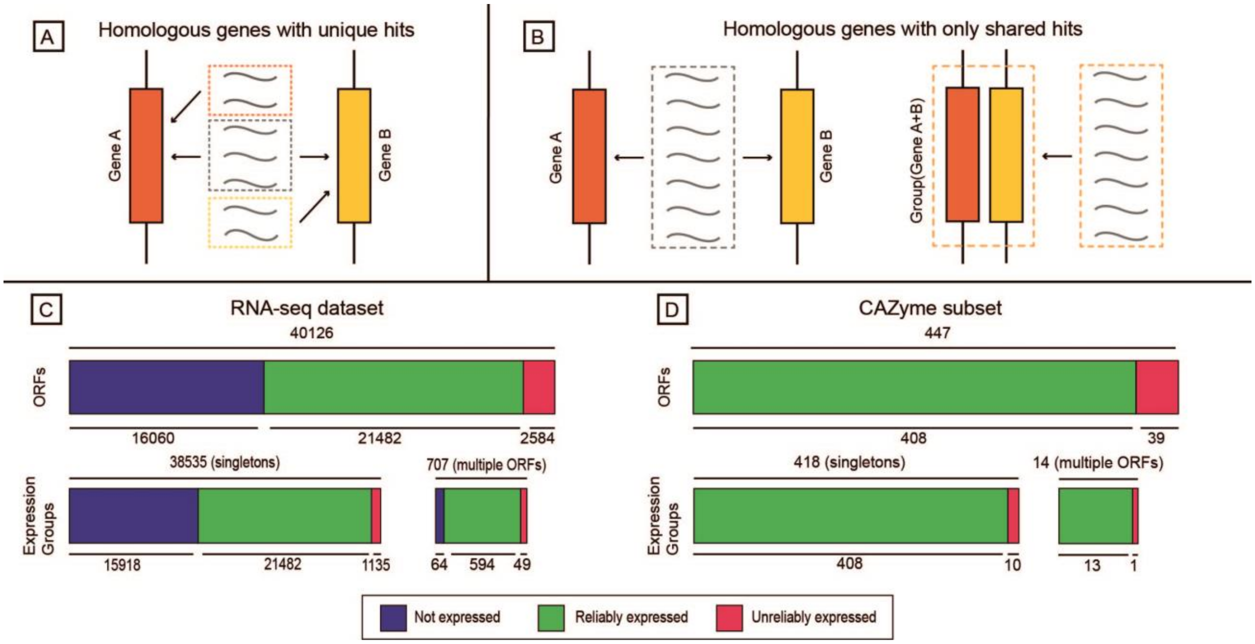


**Supplementary Figure 1. Workflow used for RNA-seq quantification and generation of expression groups.** **A** and **B** illustrate (respectively) the grouping of genes when unique reads are present (**A**) and when there are not (**B**). In **A,** genes A and B are homologous, and a set of reads maps uniquely to either gene A or to gene B. The subsequent unique sets can be used to sort the ambiguous reads that map on both genes. In **B**, genes A and B are homologous and no unique reads map exclusively to either gene. The two genes cannot be separated and thus their expression cannot be reliably estimated. Instead of losing the information, the genes are merged into an expression group. **C** and **D** represent the effect of collapsing genes into expression groups on the total dataset (**C**), and the reduced dataset focused on CAZymes (**D**).

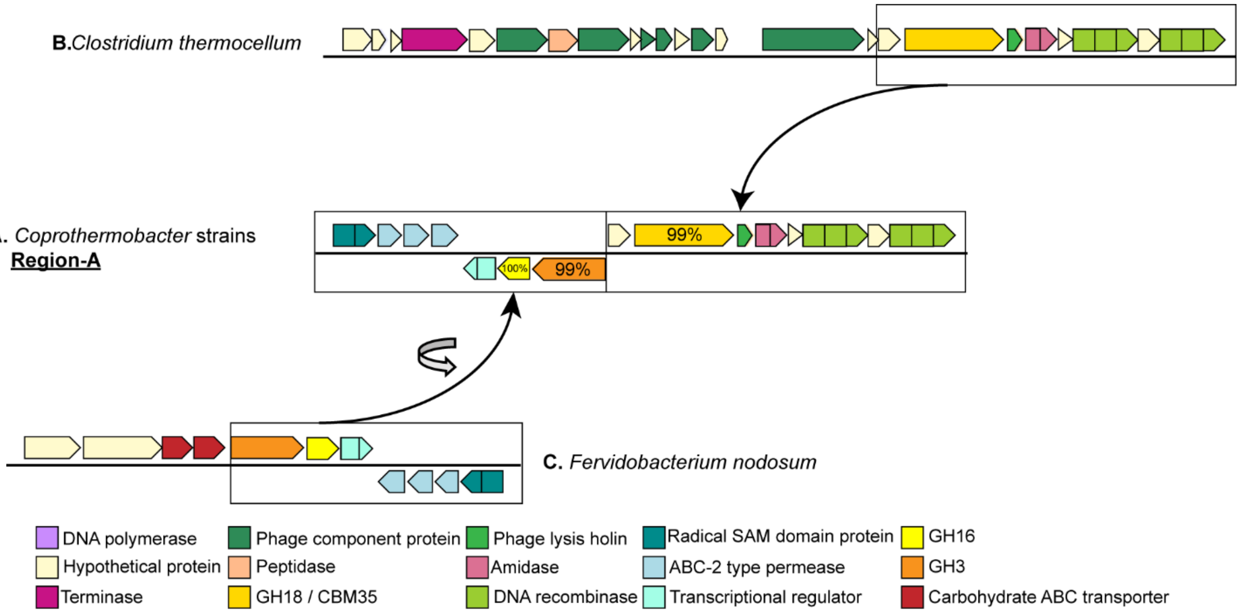


**Supplementary Figure 2. Gene synteny of CAZymes within region-A encoded in *BWF2A* and *SW3C* genomes.** **A, B** and **C** represent the gene organization of CAZymes within region-A encoded in *BWF2A* and *SW3C* (see **Figure 3**), as well as similar operons found in *C. thermocellum* and *Fervidobacterium nodosum.* Gene similarity is indicated in selected examples, and can be found in detail in **Supplementary Table 5**.

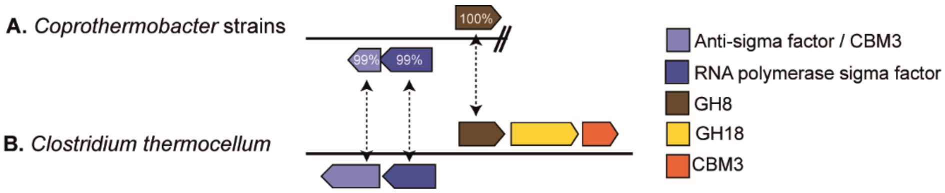


**Supplementary Figure 3. Gene organization of selected cellulosomal genes found in the *BWF2A* (A), compared against a similar operon found in *C. thermocellum* (B)*.*** Gene similarity is indicated.

**Supplementary Table 1. Anaerobic biogas reactors with reported *Coprothermobacter proteolyticus*-affiliated populations.** Date, location, reactor substrate and the presence /absence of cellulose are listed. The SEM1b consortium from this study is highlighted in yellow.

| Date | Location | Substrate | Cellulose | Hemicellulose | 16S | Reference |
|---|---|---|---|---|---|---|
| 2018 | Norway I | Cellulose (C) | YES (88.3%) | YES (9.3 %) | YES | This study |
| 2017 | Denmark | Food waste, Grass clippings (FW, C) | YES | YES | YES | (Fitamo *et al.*, 2017) |
| 2017 | Singapore | wastewater treatment plant sludge (S) | YES | n.a. | YES | (Chen *et al.*, 2017) |
| 2017 | Japan I | Acetate/sludge (Ac/S) | NO | NO | YES | (Kouzuma *et al.*, 2017) |
| 2016 | Norway II | Food waste (FW) | YES | YES | YES | (Hagen *et al.*, 2017) |
| 2016 | Korea | Seaweeds (SW) | YES | NO | YES | (Azizi *et al.*, 2016) |
| 2016 | Australia I | wastewater treatment plant sludge (S) | n.a. | n.a. | YES | (Ho *et al.*, 2016) |
| 2016 | China I | Food waste (FW) | Yes | n.a. | YES | (Chen *et al.*, 2016) |
| 2015 | Michigan | wastewater treatment plant sludge (S) | n.a. | n.a. | YES | (Sun *et al.*, 2015) |
| 2015 | China II | wastewater treatment plant sludge (S) | n.a. | n.a. | YES | (Yu *et al.*, 2015) |
| 2015 | Roma Italy | wastewater treatment plant sludge (S) | NO | NO | YES | (Gagliano *et al.*, 2015) |
| 2014 | France | Municipal Solid Waste / Office Paper (C) | YES (70%) | YES (30%) | YES | (Lü *et al.*, 2014) |
| 2013 | Australia II | wastewater treatment plant sludge (S) | n.a. | n.a. | YES | (Pervin *et al.*, 2013) |
| 2008 | Japan II | wastewater treatment plant sludge (S) | NO | NO | YES | (Kobayashi *et al.*, 2008) |

**Supplementary Table 2. 16S rRNA analysis of the SEM1b consortium.** Taxonomic composition of two separate samples (D1B and D2B), collected from a continuous SEM1b culture. D2B is 15 generations older than D1B. Consensus lineage was determined via QIIME. Closest relatives were determined using BLASTn.

| OTU ID | D1B (%) | D2B (%) | Average (%) | Consensus Lineage | Closest relative (BlastN) |
|---|---|---|---|---|---|
| OTU-1.1 | 55.3 (+/- 2.4) | 45.2 (+/- 0.2) | 50.2 | *Clostridium* | *C. thermocellum* |
| OTU-2.1 | 26.5 (+/- 1.3) | 31.4 (+/- 0.1) | 29 | *Coprothermobacter* | *C. proteolyticus* |
| OTU-2.2 | 10.6 (+/- 0.3) | 12.6 (+/- 0.1) | 11.6 | *Coprothermobacter* | *C. proteolyticus* |
| OTU-3.1 | 0 | 6.9 (+/- 0.04) | 3.5 | *Anaerobaculum* | *Acetomicrobium mobile* |
| OTU-4 | 1.5 (+/- 0.2) | 2.8 (+/- 0.1) | 2.2 | *Tepidimicrobium* | *Tepidimicrobium* |
| OTU-5 | 3.7 (+/- 1) | 0 | 1.9 | *Clostridium* | *Uncultured bacterium* |
| OTU-2.3 | 0.4 (+/- 0.1) | 0.4 (+/- 0.03) | 0.4 | *Coprothermobacter* | *C. proteolyticus* |
| OTU-3.2 | 0 | 0.6 (+/- 0) | 0.3 | *Anaerobaculum* | *Uncultured Anaerobaculum* |
| OTU-6.1 | 1.2 (+/- 0.2) | 0 | 0.6 | *Thermoanaerobacterales* | *Tepidanaerobacter* |
| OTU-6.2 | 0.7 (+/- 0.2) | 0 | 0.35 | *Thermoanaerobacterales* | *Tepidanaerobacter* |
| OTU-1.2 | 0.1 (+/- 0) | 0 | 0.05 | *Clostridium* | *C. thermocellum* |
| OTU-7 | 0.02 (+/- 0.006) | 0.02 (+/- 0.005) | 0.02 | *Methanothermobacter* | *Methanothermobacter* |

**Supplementary Table 3. CheckM and ANI analyses.** Statistics of the metagenome-recovered genomes (MAGs) from the SEM1b consortium as assessed by CheckM and average nucleotide identities (ANI), which were calculated against their closest relative.

| MAG Id | Completeness | Contamination | Heterogeneity | Size (Mb) | Closest relative | ANI (%) |
|---|---|---|---|---|---|---|
| RCLO1 | 98.7 | 0.0 | 0.0 | 3.2 | *C. thermocellum* ATCC27405 | 99.8 |
| COPR1 | 98.2 | 5.4 | 100.0 | 1.8 | *C proteolyticus* DSM5265 | 97.8 |
| COPR2 | 0.0 | 0.0 | 0.0 | 0.3 | n.a. | n.a. |
| COPR3 | 0.0 | 0.0 | 0.0 | 0.3 | n.a. | n.a. |
| SYNG1 | 86.4 | 0.0 | 0.0 | 1.4 | *Acetomicrobium mobile* DSM13181 | 98.6 |
| SYNG2 | 13.6 | 5.1 | 100.0 | 0.8 | n.a. | n.a. |
| TISS1 | 97.9 | 0.0 | 0.0 | 2.1 | *Tepidimicrobium xylanilyticum* DSM23310 | 75.1 |
| TEPI1 | 82.2 | 1.7 | 0.0 | 1.9 | *Tepidanaerobacter acetatoxydans* Re1 | 74.4 |
| TEPI2 | 87.0 | 11.8 | 14.3 | 2.4 | *Tepidanaerobacter acetatoxydans* Re1 | 74.6 |
| CLOS1 | 98.6 | 1.4 | 0.0 | 3.0 | *Clostridium Stercorarium* DSM8532 | 79.1 |
| METH1 | 99.9 | 0.0 | 0.0 | 1.7 | *Methanothermobacter thermautrophicus* | 98.2 |
| *Unbinned* | | | | 7.7 | n.a. | n.a. |

**Supplementary Table 4. CAZyme profiles for selected MAGs and genomes.** The total number of annotated domains per CAZy family for each SEM1b MAG/genome are listed. * CAZyme profile for *C. proteolyticus* DSM 5265

| CAZy family | RCLO1 | CLOS1 | TISS1 | COPR1 | COPR2 | COPR3 | *BWF2A* | *SW3C* | DSM5265* |
|---|---|---|---|---|---|---|---|---|---|
| AA3 | 0 | 0 | 2 | 0 | 0 | 0 | 0 | 0 | 0 |
| AA6 | 3 | 1 | 3 | 0 | 0 | 0 | 0 | 0 | 0 |
| CBM3 | 24 | 0 | 0 | 0 | 0 | 0 | 2 | 1 | 0 |
| CBM4 | 8 | 0 | 0 | 0 | 0 | 0 | 0 | 0 | 0 |
| CBM6 | 15 | 2 | 0 | 0 | 0 | 0 | 0 | 0 | 0 |
| CBM9 | 0 | 1 | 0 | 0 | 0 | 0 | 0 | 0 | 0 |
| CBM11 | 1 | 0 | 0 | 0 | 0 | 0 | 0 | 0 | 0 |
| CBM13 | 2 | 0 | 0 | 0 | 0 | 0 | 0 | 0 | 0 |
| CBM16 | 5 | 1 | 0 | 0 | 0 | 0 | 0 | 0 | 0 |
| CBM22 | 4 | 3 | 0 | 0 | 0 | 0 | 0 | 0 | 0 |
| CBM25 | 3 | 1 | 0 | 0 | 0 | 0 | 0 | 0 | 0 |
| CBM30 | 4 | 0 | 0 | 0 | 0 | 0 | 0 | 0 | 0 |
| CBM31 | 1 | 0 | 0 | 0 | 0 | 0 | 0 | 0 | 0 |
| CBM32 | 3 | 1 | 0 | 0 | 0 | 0 | 0 | 0 | 0 |
| CBM34 | 1 | 0 | 0 | 0 | 0 | 0 | 1 | 1 | 0 |
| CBM35 | 6 | 1 | 0 | 0 | 0 | 0 | 1 | 1 | 0 |
| CBM37 | 2 | 1 | 0 | 0 | 0 | 0 | 0 | 0 | 0 |
| CBM40 | 1 | 0 | 0 | 0 | 0 | 0 | 0 | 0 | 0 |
| CBM42 | 2 | 0 | 0 | 0 | 0 | 0 | 0 | 0 | 0 |
| CBM44 | 4 | 0 | 0 | 0 | 0 | 0 | 0 | 0 | 0 |
| CBM48 | 3 | 2 | 0 | 0 | 0 | 0 | 0 | 0 | 0 |
| CBM50 | 15 | 10 | 10 | 0 | 0 | 0 | 0 | 0 | 0 |
| CBM54 | 1 | 1 | 0 | 0 | 0 | 1 | 0 | 0 | 0 |
| CBM62 | 1 | 0 | 0 | 0 | 0 | 0 | 0 | 0 | 0 |
| CBM66 | 0 | 1 | 0 | 1 | 0 | 0 | 0 | 1 | 0 |
| CBM67 | 0 | 2 | 0 | 0 | 0 | 0 | 0 | 0 | 0 |
| CE1 | 3 | 2 | 1 | 2 | 0 | 0 | 2 | 2 | 0 |
| CE2 | 1 | 0 | 0 | 0 | 0 | 0 | 0 | 0 | 0 |
| CE3 | 2 | 1 | 0 | 0 | 0 | 0 | 0 | 0 | 0 |
| CE4 | 5 | 6 | 3 | 4 | 0 | 0 | 2 | 3 | 2 |
| CE6 | 1 | 0 | 0 | 0 | 0 | 0 | 0 | 0 | 0 |
| CE7 | 1 | 2 | 0 | 0 | 0 | 0 | 0 | 0 | 0 |
| CE8 | 1 | 1 | 0 | 0 | 0 | 0 | 0 | 0 | 0 |
| CE9 | 1 | 2 | 0 | 2 | 0 | 0 | 2 | 2 | 2 |
| CE10 | 2 | 3 | 2 | 1 | 0 | 0 | 1 | 1 | 0 |
| CE12 | 2 | 1 | 0 | 0 | 0 | 0 | 0 | 0 | 0 |
| CE14 | 1 | 0 | 0 | 1 | 0 | 0 | 1 | 1 | 0 |
| CE15 | 0 | 2 | 0 | 0 | 0 | 0 | 0 | 0 | 0 |
| cohesin | 17 | 0 | 0 | 0 | 0 | 0 | 0 | 0 | 0 |
| dockerin | 61 | 1 | 0 | 0 | 0 | 0 | 4 | 0 | 0 |
| GH1 | 2 | 1 | 0 | 0 | 0 | 0 | 0 | 0 | 0 |
| GH2 | 1 | 7 | 0 | 0 | 0 | 0 | 0 | 0 | 0 |
| GH3 | 2 | 5 | 1 | 0 | 0 | 0 | 1 | 1 | 1 |
| GH4 | 0 | 5 | 0 | 0 | 0 | 1 | 0 | 0 | 0 |

| | | | | | | | | | |
|---|---|---|---|---|---|---|---|---|---|
| **GH5** | 10 | 1 | 0 | 0 | 0 | 0 | 0 | 0 | 0 |
| **GH8** | 1 | 0 | 0 | 0 | 0 | 0 | 2 | 1 | 0 |
| **GH9** | 16 | 1 | 0 | 0 | 0 | 0 | 1 | 1 | 0 |
| **GH10** | 5 | 4 | 0 | 0 | 0 | 0 | 0 | 0 | 0 |
| **GH11** | 2 | 0 | 0 | 0 | 0 | 0 | 0 | 0 | 0 |
| **GH13** | 2 | 3 | 0 | 1 | 0 | 0 | 2 | 2 | 2 |
| **GH15** | 1 | 1 | 0 | 0 | 0 | 0 | 0 | 0 | 0 |
| **GH16** | 2 | 0 | 0 | 0 | 0 | 0 | 1 | 1 | 0 |
| **GH18** | 3 | 4 | 1 | 0 | 0 | 0 | 2 | 2 | 0 |
| **GH23** | 2 | 2 | 1 | 3 | 0 | 0 | 3 | 3 | 3 |
| **GH26** | 3 | 1 | 0 | 0 | 0 | 0 | 0 | 0 | 0 |
| **GH28** | 1 | 1 | 0 | 0 | 0 | 0 | 0 | 0 | 0 |
| **GH29** | 0 | 2 | 0 | 0 | 0 | 0 | 0 | 0 | 0 |
| **GH30** | 1 | 1 | 0 | 0 | 0 | 0 | 0 | 0 | 0 |
| **GH31** | 0 | 3 | 0 | 0 | 0 | 0 | 0 | 0 | 0 |
| **GH35** | 0 | 2 | 0 | 0 | 0 | 0 | 0 | 0 | 0 |
| **GH36** | 0 | 4 | 0 | 0 | 0 | 0 | 0 | 0 | 0 |
| **GH38** | 0 | 2 | 0 | 0 | 0 | 0 | 0 | 0 | 0 |
| **GH39** | 0 | 1 | 0 | 0 | 0 | 0 | 0 | 0 | 0 |
| **GH42** | 0 | 2 | 0 | 0 | 0 | 0 | 0 | 0 | 0 |
| **GH43** | 5 | 9 | 0 | 0 | 0 | 0 | 0 | 0 | 0 |
| **GH44** | 1 | 0 | 0 | 0 | 0 | 0 | 0 | 0 | 0 |
| **GH48** | 2 | 0 | 0 | 0 | 0 | 0 | 0 | 0 | 0 |
| **GH51** | 1 | 3 | 0 | 0 | 0 | 0 | 0 | 0 | 0 |
| **GH53** | 1 | 0 | 0 | 0 | 0 | 0 | 0 | 0 | 0 |
| **GH57** | 0 | 0 | 0 | 3 | 0 | 0 | 3 | 3 | 3 |
| **GH63** | 0 | 1 | 0 | 0 | 0 | 0 | 0 | 0 | 0 |
| **GH67** | 0 | 1 | 0 | 0 | 0 | 0 | 0 | 0 | 0 |
| **GH73** | 0 | 1 | 0 | 1 | 0 | 0 | 1 | 1 | 1 |
| **GH74** | 3 | 0 | 0 | 4 | 0 | 0 | 2 | 2 | 0 |
| **GH78** | 0 | 2 | 0 | 0 | 0 | 0 | 0 | 0 | 0 |
| **GH81** | 1 | 0 | 0 | 0 | 0 | 0 | 0 | 0 | 0 |
| **GH88** | 0 | 2 | 0 | 0 | 0 | 0 | 0 | 0 | 0 |
| **GH94** | 3 | 1 | 1 | 0 | 0 | 0 | 0 | 0 | 0 |
| **GH95** | 0 | 5 | 0 | 0 | 0 | 0 | 0 | 0 | 0 |
| **GH105** | 0 | 6 | 0 | 0 | 0 | 0 | 0 | 0 | 0 |
| **GH106** | 0 | 3 | 0 | 0 | 0 | 0 | 0 | 0 | 0 |
| **GH109** | 2 | 10 | 4 | 7 | 3 | 1 | 3 | 3 | 0 |
| **GH112** | 0 | 1 | 0 | 0 | 0 | 0 | 0 | 0 | 0 |
| **GH115** | 0 | 1 | 0 | 0 | 0 | 0 | 0 | 0 | 0 |
| **GH120** | 0 | 1 | 0 | 0 | 0 | 0 | 0 | 0 | 0 |
| **GH124** | 3 | 0 | 0 | 0 | 0 | 0 | 0 | 0 | 0 |
| **GH126** | 1 | 0 | 0 | 0 | 0 | 0 | 0 | 0 | 0 |
| **GH127** | 0 | 1 | 0 | 0 | 0 | 0 | 0 | 0 | 0 |
| **GH130** | 1 | 4 | 0 | 0 | 0 | 0 | 0 | 0 | 0 |
| **GH133** | 1 | 1 | 0 | 0 | 0 | 0 | 0 | 0 | 0 |
| **PL1** | 2 | 0 | 0 | 0 | 0 | 0 | 0 | 0 | 0 |
| **PL9** | 1 | 1 | 0 | 0 | 0 | 0 | 0 | 0 | 0 |

| | | | | | | | | | |
|---|---|---|---|---|---|---|---|---|---|
| **PL11** | 1 | 1 | 0 | 0 | 0 | 0 | 0 | 0 | 0 |
| **PL12** | 0 | 2 | 0 | 0 | 0 | 2 | 0 | 0 | 0 |
| **PL22** | 0 | 1 | 0 | 0 | 0 | 0 | 0 | 0 | 0 |
| Total | 290 | 160 | 29 | 30 | 3 | 5 | 37 | 33 | 14 |

**Supplementary Table 5. Selected CAZymes that were acquired within BWF2A and SW3C.** The CAZymes are listed for the two strains (IMG genome ID: 2731957509 and 2731957514, respectively) with their respective Gene IMG IDs. CAZymes found in region-A (**Figure 3**) are shaded in purple. CAZy family assignment and similarity % with their closest relative are indicated.

| Gene IMG ID (BWF2A) | CAZy family | Best hit and similarity (%) | Gene ID (SW3C) | CAZy family | Best hit and similarity (%) |
|---|---|---|---|---|---|
| 2731988480_Ga0187557_1002248 | GH16 | *F. nodosum* (100) | 2731996536_Ga0187564_10674 | GH16 | *F. nodosum* (100) |
| 2731988481_Ga0187557_1002249 | GH3 | *F. nodosum* (99) | 2731996537_Ga0187564_10675 | GH3 | *F. nodosum* (99) |
| 2731988483_Ga0187557_1002251 | GH18-CBM35 | *C. thermocellum* (99) | 2731996539_Ga0187564_10677 | GH18-CBM35 | *C. thermocellum* (99) |
| 2731989281_Ga0187557_10084 | GH8 | *C. thermocellum* (99) | 2731996671_Ga0187564_1131 | GH36 | *C. thermocellum* (98) |
| 2731989282_Ga0187557_10091 | GH9-doc | *C. thermocellum* (99) | 2731996673_Ga0187564_1133 | GH9 | *C. thermocellum* (100) |
| 2731989288_Ga0187557_10111 | GH36 | *C. thermocellum* (100) | 2731996687_Ga0187564_1201 | GH8 | *C. thermocellum* (99) |
| 2731989293_Ga0187557_10122 | CBM3 | *C. thermocellum* (100) | 2731996693_Ga0187564_1232 | GH18-doc | *C. thermocellum* (99) |
| 2731989294_Ga0187557_10123 | GH18-doc | *C. thermocellum* (96) | 2731996702_Ga0187564_1292 | CBM3 | *C. thermocellum* (100) |
| 2731989310_Ga0187557_10201 | CBM3 | *C. thermocellum* (100) | | | |
| 2731989313_Ga0187557_10212 | GH8-doc | *C. thermocellum* (100) | | | |

**Supplementary Table 6. Clusters of expression groups.** The table shows the gene IMG ID and the CAZy family assignment of each expression group together with the genome and cluster. Groups composed of several ORFs contain a "+" between the gene IDs and the names of the organisms. CAZymes in expression groups without uniquely mapping reads (unique hits) are not part of the clustering and were labelled as "n.a.".

| Expression Groups (Gene IMG ID) | CAZY | Genome | Cluster |
|---|---|---|---|
| Ga0196617_10001933 | CBM25 | RCLO1 | I |
| Ga0196617_100013115 | GH109 | TISS1 | I |
| Ga0196617_1002411 | GH109 | COPR2 | I |
| Ga0196617_1002669 | GH4 | COPR3 | I |
| Ga0187557_1002157 | GH109 | BWF2A | I |
| Ga0187557_1003280 | GH73 | BWF2A | I |
| Ga0187564_102294 | GH109 | SW3C | I |
| Ga0187564_103104 | CE14 | SW3C | I |
| Ga0187564_104108 | GH23 | SW3C | I |
| Ga0187564_10645 | GH109 | SW3C | I |
| Ga0187564_10677 | CBM35 | SW3C | I |
| Ga0187564_10677 | GH18 | SW3C | I |
| Ga0187557_100270+Ga0187564_103120 | CE9 | SW3C + BWF2A | I |
| Ga0196617_1000499 | GH23 | COPR1 | II |
| Ga0196617_1000628 | CE9 | COPR1 | II |
| Ga0196617_10022612 | GH109 | COPR1 | II |
| Ga0196617_1003256 | CE1 | COPR1 | II |
| Ga0196617_1006634 | GH109 | COPR1 | II |
| Ga0196617_1009191 | GH109 | COPR1 | II |
| Ga0187557_1002223 | GH109 | BWF2A | II |
| Ga0187564_10257 | SLH | SW3C | II |
| Ga0187564_10257 | SLH | SW3C | II |
| Ga0187564_1201 | GH8 | SW3C | II |
| Ga0187557_1002314+Ga0187564_106139 | CBM34 | SW3C + BWF2A | II |
| Ga0187557_1002314+Ga0187564_106139 | GH13 | SW3C + BWF2A | II |
| Ga0187564_101146+Ga0187557_1001161 | GH23 | SW3C + BWF2A | II |
| Ga0196617_1000454 | CBM6 | RCLO1 | III |
| Ga0196617_1000454 | CE1 | RCLO1 | III |
| Ga0196617_1000454 | dockerin | RCLO1 | III |
| Ga0196617_10009713 | CBM50 | RCLO1 | III |
| Ga0196617_10014620 | CE4 | COPR1 | III |
| Ga0196617_1004347 | GH74 | COPR1 | III |
| Ga0196617_1005644 | GH109 | COPR1 | III |
| Ga0196617_10005857 | AA6 | TISS1 | III |
| Ga0187557_100358 | SLH | BWF2A | III |
| Ga0187557_100358 | SLH | BWF2A | III |
| Ga0187564_10258 | SLH | SW3C | III |
| Ga0187564_10258 | SLH | SW3C | III |
| Ga0187564_103147 | GH57 | SW3C | III |
| Ga0187564_10471 | GH13 | SW3C | III |
| Ga0187564_10493 | CE4 | SW3C | III |
| Ga0196617_100009115 | CBM13 | RCLO1 | IV |

| | | | |
|---|---|---|---|
| Ga0196617_100009115 | GH43 | RCLO1 | IV |
| Ga0196617_100009116 | dockerin | RCLO1 | IV |
| Ga0196617_100009116 | dockerin | RCLO1 | IV |
| Ga0196617_100009116 | GH81 | RCLO1 | IV |
| Ga0196617_100009143 | CBM48 | RCLO1 | IV |
| Ga0196617_100009158 | CBM3 | RCLO1 | IV |
| Ga0196617_100009158 | GH9 | RCLO1 | IV |
| Ga0196617_100009159 | CBM30 | RCLO1 | IV |
| Ga0196617_100009159 | CBM44 | RCLO1 | IV |
| Ga0196617_100009159 | GH44 | RCLO1 | IV |
| Ga0196617_100009159 | GH9 | RCLO1 | IV |
| Ga0196617_100009203 | CBM3 | RCLO1 | IV |
| Ga0196617_100009203 | GH9 | RCLO1 | IV |
| Ga0196617_100009224 | CE4 | RCLO1 | IV |
| Ga0196617_10000923 | CBM3 | RCLO1 | IV |
| Ga0196617_10000923 | dockerin | RCLO1 | IV |
| Ga0196617_10000923 | GH9 | RCLO1 | IV |
| Ga0196617_10000935 | cohesin | RCLO1 | IV |
| Ga0196617_10000935 | cohesin | RCLO1 | IV |
| Ga0196617_10000935 | cohesin | RCLO1 | IV |
| Ga0196617_10000935 | cohesin | RCLO1 | IV |
| Ga0196617_10000935 | cohesin | RCLO1 | IV |
| Ga0196617_10000935 | cohesin | RCLO1 | IV |
| Ga0196617_10000935 | cohesin | RCLO1 | IV |
| Ga0196617_10000936 | cohesin | RCLO1 | IV |
| Ga0196617_100014143 | CBM30 | RCLO1 | IV |
| Ga0196617_100014143 | GH9 | RCLO1 | IV |
| Ga0196617_100014145 | CBM32 | RCLO1 | IV |
| Ga0196617_100014145 | dockerin | RCLO1 | IV |
| Ga0196617_100014145 | GH5 | RCLO1 | IV |
| Ga0196617_10001446 | dockerin | RCLO1 | IV |
| Ga0196617_10001449 | CBM32 | RCLO1 | IV |
| Ga0196617_10001451 | CBM22 | RCLO1 | IV |
| Ga0196617_10001451 | CBM22 | RCLO1 | IV |
| Ga0196617_10001451 | CE1 | RCLO1 | IV |
| Ga0196617_10001451 | GH10 | RCLO1 | IV |
| Ga0196617_1000146 | CBM25 | RCLO1 | IV |
| Ga0196617_10001484 | GH23 | RCLO1 | IV |
| Ga0196617_10001497 | AA6 | RCLO1 | IV |
| Ga0196617_10001610 | CBM3 | RCLO1 | IV |
| Ga0196617_10001610 | CBM40 | RCLO1 | IV |
| Ga0196617_10001610 | SLH | RCLO1 | IV |
| Ga0196617_10001610 | SLH | RCLO1 | IV |
| Ga0196617_10001610 | SLH | RCLO1 | IV |
| Ga0196617_100016105 | SLH | RCLO1 | IV |
| Ga0196617_100016107 | SLH | RCLO1 | IV |
| Ga0196617_100016107 | SLH | RCLO1 | IV |
| Ga0196617_1000162 | CBM37 | RCLO1 | IV |

| | | | |
|---|---|---|---|
| Ga0196617_1000162 | SLH | RCLO1 | IV |
| Ga0196617_1000162 | SLH | RCLO1 | IV |
| Ga0196617_1000162 | SLH | RCLO1 | IV |
| Ga0196617_10001654 | GH51 | RCLO1 | IV |
| Ga0196617_10001684 | CBM22 | RCLO1 | IV |
| Ga0196617_10001684 | dockerin | RCLO1 | IV |
| Ga0196617_10001684 | dockerin | RCLO1 | IV |
| Ga0196617_10001684 | GH10 | RCLO1 | IV |
| Ga0196617_100019129 | CBM31 | RCLO1 | IV |
| Ga0196617_10001939 | CE4 | RCLO1 | IV |
| Ga0196617_100023113 | GH109 | RCLO1 | IV |
| Ga0196617_10002318 | CBM11 | RCLO1 | IV |
| Ga0196617_10002318 | dockerin | RCLO1 | IV |
| Ga0196617_10002318 | dockerin | RCLO1 | IV |
| Ga0196617_10002318 | GH26 | RCLO1 | IV |
| Ga0196617_10002318 | GH5 | RCLO1 | IV |
| Ga0196617_10002319 | GH5 | RCLO1 | IV |
| Ga0196617_10002366 | GH1 | RCLO1 | IV |
| Ga0196617_10002394 | GH53 | RCLO1 | IV |
| Ga0196617_10002396 | GH74 | RCLO1 | IV |
| Ga0196617_10002396 | GH74 | RCLO1 | IV |
| Ga0196617_10002396 | GH74 | RCLO1 | IV |
| Ga0196617_100024103 | CBM3 | RCLO1 | IV |
| Ga0196617_100024106 | SLH | RCLO1 | IV |
| Ga0196617_100024106 | SLH | RCLO1 | IV |
| Ga0196617_100024106 | SLH | RCLO1 | IV |
| Ga0196617_100024109 | SLH | RCLO1 | IV |
| Ga0196617_100024109 | SLH | RCLO1 | IV |
| Ga0196617_100024109 | SLH | RCLO1 | IV |
| Ga0196617_10002441 | GH133 | RCLO1 | IV |
| Ga0196617_10002447 | dockerin | RCLO1 | IV |
| Ga0196617_10002490 | CBM3 | RCLO1 | IV |
| Ga0196617_10002490 | GH48 | RCLO1 | IV |
| Ga0196617_100025109 | SLH | RCLO1 | IV |
| Ga0196617_10002537 | dockerin | RCLO1 | IV |
| Ga0196617_10002537 | dockerin | RCLO1 | IV |
| Ga0196617_10002556 | CBM37 | RCLO1 | IV |
| Ga0196617_100026113 | CBM4 | RCLO1 | IV |
| Ga0196617_100026113 | dockerin | RCLO1 | IV |
| Ga0196617_100026113 | GH28 | RCLO1 | IV |
| Ga0196617_10002620 | GH10 | RCLO1 | IV |
| Ga0196617_10002651 | dockerin | RCLO1 | IV |
| Ga0196617_10002651 | dockerin | RCLO1 | IV |
| Ga0196617_10002651 | GH48 | RCLO1 | IV |
| Ga0196617_10003331 | CBM50 | RCLO1 | IV |
| Ga0196617_10003331 | CBM50 | RCLO1 | IV |
| Ga0196617_10003331 | GH18 | RCLO1 | IV |
| Ga0196617_10003345 | GH15 | RCLO1 | IV |

| | | | |
|---|---|---|---|
| Ga0196617_100035100 | CBM30 | RCLO1 | IV |
| Ga0196617_100035100 | CBM3 | RCLO1 | IV |
| Ga0196617_100035100 | CBM4 | RCLO1 | IV |
| Ga0196617_100035100 | dockerin | RCLO1 | IV |
| Ga0196617_100035100 | GH9 | RCLO1 | IV |
| Ga0196617_100035101 | dockerin | RCLO1 | IV |
| Ga0196617_100035101 | GH9 | RCLO1 | IV |
| Ga0196617_10003560 | cohesin | RCLO1 | IV |
| Ga0196617_10003575 | dockerin | RCLO1 | IV |
| Ga0196617_10003578 | GH124 | RCLO1 | IV |
| Ga0196617_10003580 | CBM3 | RCLO1 | IV |
| Ga0196617_10003580 | GH9 | RCLO1 | IV |
| Ga0196617_10003947 | SLH | RCLO1 | IV |
| Ga0196617_10003947 | SLH | RCLO1 | IV |
| Ga0196617_10003947 | SLH | RCLO1 | IV |
| Ga0196617_10003956 | GH3 | RCLO1 | IV |
| Ga0196617_10003979 | CBM50 | RCLO1 | IV |
| Ga0196617_10004017 | GH94 | RCLO1 | IV |
| Ga0196617_10004018 | dockerin | RCLO1 | IV |
| Ga0196617_10004018 | dockerin | RCLO1 | IV |
| Ga0196617_10004018 | GH9 | RCLO1 | IV |
| Ga0196617_10004022 | GH18 | RCLO1 | IV |
| Ga0196617_10004023 | dockerin | RCLO1 | IV |
| Ga0196617_10004023 | dockerin | RCLO1 | IV |
| Ga0196617_10004023 | GH8 | RCLO1 | IV |
| Ga0196617_10004025 | CBM3 | RCLO1 | IV |
| Ga0196617_10004046 | CBM35 | RCLO1 | IV |
| Ga0196617_10004046 | PL11 | RCLO1 | IV |
| Ga0196617_10004053 | dockerin | RCLO1 | IV |
| Ga0196617_10004053 | dockerin | RCLO1 | IV |
| Ga0196617_10004125 | CBM25 | RCLO1 | IV |
| Ga0196617_10004148 | CBM35 | RCLO1 | IV |
| Ga0196617_10004148 | CE12 | RCLO1 | IV |
| Ga0196617_10004149 | CE12 | RCLO1 | IV |
| Ga0196617_10004149 | dockerin | RCLO1 | IV |
| Ga0196617_10004156 | dockerin | RCLO1 | IV |
| Ga0196617_10004156 | dockerin | RCLO1 | IV |
| Ga0196617_10004167 | SLH | RCLO1 | IV |
| Ga0196617_10004167 | SLH | RCLO1 | IV |
| Ga0196617_10004167 | SLH | RCLO1 | IV |
| Ga0196617_10004178 | GH130 | RCLO1 | IV |
| Ga0196617_1000447 | SLH | RCLO1 | IV |
| Ga0196617_1000447 | SLH | RCLO1 | IV |
| Ga0196617_1000447 | SLH | RCLO1 | IV |
| Ga0196617_10004476 | cohesin | RCLO1 | IV |
| Ga0196617_10004476 | SLH | RCLO1 | IV |
| Ga0196617_10004476 | SLH | RCLO1 | IV |
| Ga0196617_10004476 | SLH | RCLO1 | IV |

| | | | |
|---|---|---|---|
| Ga0196617_1000451 | CBM6 | RCLO1 | IV |
| Ga0196617_1000451 | dockerin | RCLO1 | IV |
| Ga0196617_1000451 | dockerin | RCLO1 | IV |
| Ga0196617_1000452 | CBM6 | RCLO1 | IV |
| Ga0196617_1000452 | GH43 | RCLO1 | IV |
| Ga0196617_10004521 | CBM35 | RCLO1 | IV |
| Ga0196617_10004521 | GH124 | RCLO1 | IV |
| Ga0196617_10004521 | PL1 | RCLO1 | IV |
| Ga0196617_10004521 | PL9 | RCLO1 | IV |
| Ga0196617_1000453 | CBM6 | RCLO1 | IV |
| Ga0196617_1000453 | dockerin | RCLO1 | IV |
| Ga0196617_1000453 | dockerin | RCLO1 | IV |
| Ga0196617_10004533 | GH126 | RCLO1 | IV |
| Ga0196617_10004546 | SLH | RCLO1 | IV |
| Ga0196617_10004546 | SLH | RCLO1 | IV |
| Ga0196617_10004546 | SLH | RCLO1 | IV |
| Ga0196617_1000455 | CBM6 | RCLO1 | IV |
| Ga0196617_1000455 | dockerin | RCLO1 | IV |
| Ga0196617_1000455 | dockerin | RCLO1 | IV |
| Ga0196617_1000455 | GH43 | RCLO1 | IV |
| Ga0196617_10004552 | CBM16 | RCLO1 | IV |
| Ga0196617_10004553 | CBM3 | RCLO1 | IV |
| Ga0196617_10004553 | GH5 | RCLO1 | IV |
| Ga0196617_1000456 | CBM13 | RCLO1 | IV |
| Ga0196617_1000456 | CBM62 | RCLO1 | IV |
| Ga0196617_1000456 | CBM6 | RCLO1 | IV |
| Ga0196617_1000456 | dockerin | RCLO1 | IV |
| Ga0196617_1000456 | dockerin | RCLO1 | IV |
| Ga0196617_1000456 | GH5 | RCLO1 | IV |
| Ga0196617_1000458 | CBM48 | RCLO1 | IV |
| Ga0196617_1000458 | CBM48 | RCLO1 | IV |
| Ga0196617_1000458 | GH13 | RCLO1 | IV |
| Ga0196617_1000459 | CE9 | RCLO1 | IV |
| Ga0196617_10004620 | GH3 | RCLO1 | IV |
| Ga0196617_10004621 | CBM3 | RCLO1 | IV |
| Ga0196617_10004621 | CBM4 | RCLO1 | IV |
| Ga0196617_10004635 | CBM6 | RCLO1 | IV |
| Ga0196617_10004635 | CBM6 | RCLO1 | IV |
| Ga0196617_10004635 | dockerin | RCLO1 | IV |
| Ga0196617_10004635 | dockerin | RCLO1 | IV |
| Ga0196617_10004635 | GH43 | RCLO1 | IV |
| Ga0196617_10004637 | CBM42 | RCLO1 | IV |
| Ga0196617_10006312 | CBM16 | RCLO1 | IV |
| Ga0196617_10006312 | CBM16 | RCLO1 | IV |
| Ga0196617_10006336 | SLH | RCLO1 | IV |
| Ga0196617_1000635 | dockerin | RCLO1 | IV |
| Ga0196617_1000635 | GH9 | RCLO1 | IV |
| Ga0196617_10006355 | CBM3 | RCLO1 | IV |

| | | | |
|---|---|---|---|
| Ga0196617_10006355 | GH9 | RCLO1 | IV |
| Ga0196617_10006356 | CBM3 | RCLO1 | IV |
| Ga0196617_10006356 | CBM3 | RCLO1 | IV |
| Ga0196617_10006356 | dockerin | RCLO1 | IV |
| Ga0196617_10006356 | dockerin | RCLO1 | IV |
| Ga0196617_10006356 | GH9 | RCLO1 | IV |
| Ga0196617_1000636 | CBM35 | RCLO1 | IV |
| Ga0196617_1000636 | GH26 | RCLO1 | IV |
| Ga0196617_1000731 | CBM44 | RCLO1 | IV |
| Ga0196617_1000731 | CBM44 | RCLO1 | IV |
| Ga0196617_1000731 | SLH | RCLO1 | IV |
| Ga0196617_1000731 | SLH | RCLO1 | IV |
| Ga0196617_1000731 | SLH | RCLO1 | IV |
| Ga0196617_10007916 | CBM6 | RCLO1 | IV |
| Ga0196617_10007916 | CE1 | RCLO1 | IV |
| Ga0196617_10007916 | GH10 | RCLO1 | IV |
| Ga0196617_10007950 | SLH | RCLO1 | IV |
| Ga0196617_10007951 | SLH | RCLO1 | IV |
| Ga0196617_10007951 | SLH | RCLO1 | IV |
| Ga0196617_10008049 | CBM6 | RCLO1 | IV |
| Ga0196617_10008049 | GH30 | RCLO1 | IV |
| Ga0196617_1000831 | CBM44 | RCLO1 | IV |
| Ga0196617_1000831 | GH124 | RCLO1 | IV |
| Ga0196617_10008313 | CE4 | RCLO1 | IV |
| Ga0196617_10008321 | CBM6 | RCLO1 | IV |
| Ga0196617_10008321 | CBM6 | RCLO1 | IV |
| Ga0196617_10008321 | SLH | RCLO1 | IV |
| Ga0196617_10008321 | SLH | RCLO1 | IV |
| Ga0196617_1000954 | CBM22 | RCLO1 | IV |
| Ga0196617_1000954 | dockerin | RCLO1 | IV |
| Ga0196617_1000954 | dockerin | RCLO1 | IV |
| Ga0196617_1000954 | GH10 | RCLO1 | IV |
| Ga0196617_1000961 | CBM30 | RCLO1 | IV |
| Ga0196617_1000961 | CBM4 | RCLO1 | IV |
| Ga0196617_10009628 | dockerin | RCLO1 | IV |
| Ga0196617_1000968 | dockerin | RCLO1 | IV |
| Ga0196617_1000968 | dockerin | RCLO1 | IV |
| Ga0196617_1000968 | GH5 | RCLO1 | IV |
| Ga0196617_1000969 | CBM3 | RCLO1 | IV |
| Ga0196617_10009710 | SLH | RCLO1 | IV |
| Ga0196617_10009710 | SLH | RCLO1 | IV |
| Ga0196617_10009729 | CBM50 | RCLO1 | IV |
| Ga0196617_10010619 | SLH | RCLO1 | IV |
| Ga0196617_10010619 | SLH | RCLO1 | IV |
| Ga0196617_10010641 | AA6 | RCLO1 | IV |
| Ga0196617_10010817 | GH23 | RCLO1 | IV |
| Ga0196617_1001211 | CBM42 | RCLO1 | IV |
| Ga0196617_1001211 | GH43 | RCLO1 | IV |

| | | | |
|---|---|---|---|
| Ga0196617_10012123 | CBM35 | RCLO1 | IV |
| Ga0196617_10012123 | dockerin | RCLO1 | IV |
| Ga0196617_10012123 | dockerin | RCLO1 | IV |
| Ga0196617_10012123 | GH26 | RCLO1 | IV |
| Ga0196617_10012132 | CBM3 | RCLO1 | IV |
| Ga0196617_10012132 | CBM3 | RCLO1 | IV |
| Ga0196617_10012132 | GH9 | RCLO1 | IV |
| Ga0196617_10012135 | CBM3 | RCLO1 | IV |
| Ga0196617_10012135 | dockerin | RCLO1 | IV |
| Ga0196617_10012135 | dockerin | RCLO1 | IV |
| Ga0196617_10012135 | GH9 | RCLO1 | IV |
| Ga0196617_10012136 | dockerin | RCLO1 | IV |
| Ga0196617_10012136 | dockerin | RCLO1 | IV |
| Ga0196617_10012516 | CBM3 | RCLO1 | IV |
| Ga0196617_10012516 | dockerin | RCLO1 | IV |
| Ga0196617_10012516 | dockerin | RCLO1 | IV |
| Ga0196617_10012516 | GH9 | RCLO1 | IV |
| Ga0196617_1001259 | dockerin | RCLO1 | IV |
| Ga0196617_1001259 | dockerin | RCLO1 | IV |
| Ga0196617_1001259 | GH5 | RCLO1 | IV |
| Ga0196617_10012616 | GH94 | RCLO1 | IV |
| Ga0196617_10012633 | CBM50 | RCLO1 | IV |
| Ga0196617_10012633 | CBM50 | RCLO1 | IV |
| Ga0196617_10012633 | CBM50 | RCLO1 | IV |
| Ga0196617_10012633 | CBM50 | RCLO1 | IV |
| Ga0196617_10012634 | CBM50 | RCLO1 | IV |
| Ga0196617_10012634 | CBM50 | RCLO1 | IV |
| Ga0196617_10012634 | CBM50 | RCLO1 | IV |
| Ga0196617_10012635 | CBM50 | RCLO1 | IV |
| Ga0196617_10012635 | CBM50 | RCLO1 | IV |
| Ga0196617_10012711 | SLH | RCLO1 | IV |
| Ga0196617_10012711 | SLH | RCLO1 | IV |
| Ga0196617_10012711 | SLH | RCLO1 | IV |
| Ga0196617_10012713 | CE6 | RCLO1 | IV |
| Ga0196617_10012729 | GH18 | RCLO1 | IV |
| Ga0196617_1001275 | GH5 | RCLO1 | IV |
| Ga0196617_10012833 | CBM3 | RCLO1 | IV |
| Ga0196617_1001361 | CE3 | RCLO1 | IV |
| Ga0196617_1001361 | CE3 | RCLO1 | IV |
| Ga0196617_1001361 | dockerin | RCLO1 | IV |
| Ga0196617_1001361 | dockerin | RCLO1 | IV |
| Ga0196617_10013629 | CBM50 | RCLO1 | IV |
| Ga0196617_1001363 | CE2 | RCLO1 | IV |
| Ga0196617_1001363 | GH5 | RCLO1 | IV |
| Ga0196617_1001365 | CBM34 | RCLO1 | IV |
| Ga0196617_1001365 | GH13 | RCLO1 | IV |
| Ga0196617_10013919 | GH16 | RCLO1 | IV |
| Ga0196617_10013921 | GH1 | RCLO1 | IV |

| | | | |
|---|---|---|---|
| Ga0196617_10014113 | CE10 | RCLO1 | IV |
| Ga0196617_10014123 | CE14 | RCLO1 | IV |
| Ga0196617_1001491 | cohesin | RCLO1 | IV |
| Ga0196617_1001491 | cohesin | RCLO1 | IV |
| Ga0196617_1001491 | SLH | RCLO1 | IV |
| Ga0196617_1001491 | SLH | RCLO1 | IV |
| Ga0196617_1001491 | SLH | RCLO1 | IV |
| Ga0196617_1001492 | cohesin | RCLO1 | IV |
| Ga0196617_1001492 | SLH | RCLO1 | IV |
| Ga0196617_1001492 | SLH | RCLO1 | IV |
| Ga0196617_1001492 | SLH | RCLO1 | IV |
| Ga0196617_10014924 | CBM16 | RCLO1 | IV |
| Ga0196617_10015118 | CE4 | RCLO1 | IV |
| Ga0196617_10019112 | GH109 | RCLO1 | IV |
| Ga0196617_10019617 | CBM3 | RCLO1 | IV |
| Ga0196617_10019617 | CBM3 | RCLO1 | IV |
| Ga0196617_10019617 | GH9 | RCLO1 | IV |
| Ga0196617_1002189 | CBM35 | RCLO1 | IV |
| Ga0196617_1002189 | PL1 | RCLO1 | IV |
| Ga0196617_1002401 | CBM3 | RCLO1 | IV |
| Ga0196617_1002401 | cohesin | RCLO1 | IV |
| Ga0196617_1002401 | cohesin | RCLO1 | IV |
| Ga0196617_1002401 | cohesin | RCLO1 | IV |
| Ga0196617_1002401 | cohesin | RCLO1 | IV |
| Ga0196617_10024516 | CE4 | RCLO1 | IV |
| Ga0196617_10024517 | CBM6 | RCLO1 | IV |
| Ga0196617_10024517 | dockerin | RCLO1 | IV |
| Ga0196617_10024517 | GH11 | RCLO1 | IV |
| Ga0196617_10024518 | CBM6 | RCLO1 | IV |
| Ga0196617_10024518 | dockerin | RCLO1 | IV |
| Ga0196617_10024518 | dockerin | RCLO1 | IV |
| Ga0196617_10024518 | GH11 | RCLO1 | IV |
| Ga0196617_1002738 | CE10 | RCLO1 | IV |
| Ga0196617_1005678 | CE7 | RCLO1 | IV |
| Ga0196617_1008234 | GH109 | COPR1 | IV |
| Ga0196617_100004299 | AA6 | TISS1 | IV |
| Ga0196617_100012177 | CBM50 | TISS1 | IV |
| Ga0196617_1000123 | AA6 | TISS1 | IV |
| Ga0196617_100005248 | CBM22 | CLOS1 | IV |
| Ga0196617_100005248 | CBM22 | CLOS1 | IV |
| Ga0196617_100005248 | CBM9 | CLOS1 | IV |
| Ga0196617_100005248 | GH10 | CLOS1 | IV |
| Ga0196617_10000543 | GH130 | CLOS1 | IV |
| Ga0196617_1006694 | GH115 | CLOS1 | IV |
| Ga0196617_1008072 | GH43 | CLOS1 | IV |
| Ga0196617_1010211 | GH4 | CLOS1 | IV |
| Ga0196617_1004561 | GH109 | COPR2 | IV |
| Ga0196617_1002048 | CBM54 | COPR3 | IV |

| | | | |
|---|---|---|---|
| Ga0196617_1002048 | SLH | COPR3 | IV |
| Ga0196617_1002048 | SLH | COPR3 | IV |
| Ga0196617_1002048 | SLH | COPR3 | IV |
| Ga0196617_1002716 | GH109 | COPR3 | IV |
| Ga0196617_10004021+Ga0187557_10122 | CBM3 | BWF2A + CLOS1 | IV |
| Ga0187564_1232 | GH18 | SW3C | IV |
| Ga0196617_1005697 | GH73 | COPR1 | V |
| Ga0196617_100010102 | GH109 | TISS1 | V |
| Ga0196617_10001093 | GH109 | TISS1 | V |
| Ga0196617_10001250 | CBM50 | TISS1 | V |
| Ga0196617_10001250 | CBM50 | TISS1 | V |
| Ga0196617_10001250 | CBM50 | TISS1 | V |
| Ga0196617_10000516 | GH2 | CLOS1 | V |
| Ga0196617_100005270 | SLH | CLOS1 | V |
| Ga0196617_10001198 | SLH | CLOS1 | V |
| Ga0196617_10001198 | SLH | CLOS1 | V |
| Ga0196617_100018135 | CE4 | CLOS1 | V |
| Ga0196617_10003852 | GH95 | CLOS1 | V |
| Ga0196617_10012046 | GH109 | CLOS1 | V |
| Ga0187564_102193 | CE1 | SW3C | V |
| Ga0187564_10318 | GH109 | SW3C | V |
| Ga0187564_10322 | CE9 | SW3C | V |
| Ga0196617_10006310 | GH5 | RCLO1 | VI |
| Ga0196617_1000638 | CBM4 | RCLO1 | VI |
| Ga0196617_1000638 | CBM4 | RCLO1 | VI |
| Ga0196617_1000638 | CBM4 | RCLO1 | VI |
| Ga0196617_1000638 | CBM4 | RCLO1 | VI |
| Ga0196617_1000638 | CBM54 | RCLO1 | VI |
| Ga0196617_1000638 | GH16 | RCLO1 | VI |
| Ga0196617_1000638 | SLH | RCLO1 | VI |
| Ga0196617_1000638 | SLH | RCLO1 | VI |
| Ga0196617_1000638 | SLH | RCLO1 | VI |
| Ga0196617_10006233 | GH57 | COPR1 | VI |
| Ga0196617_10006234 | GH57 | COPR1 | VI |
| Ga0196617_10006619 | GH23 | COPR1 | VI |
| Ga0196617_10006658 | GH13 | COPR1 | VI |
| Ga0196617_1001466 | GH23 | COPR1 | VI |
| Ga0196617_10018610 | CE14 | COPR1 | VI |
| Ga0196617_1002859 | GH74 | COPR1 | VI |
| Ga0196617_10031312 | CE4 | COPR1 | VI |
| Ga0196617_1003146 | SLH | COPR1 | VI |
| Ga0196617_1003797 | GH109 | COPR1 | VI |
| Ga0196617_1003936 | CE1 | COPR1 | VI |
| Ga0196617_1006193 | SLH | COPR1 | VI |
| Ga0196617_1006193 | SLH | COPR1 | VI |
| Ga0196617_1006193 | SLH | COPR1 | VI |
| Ga0196617_1006445 | GH109 | COPR1 | VI |
| Ga0196617_1009463 | CE9 | COPR1 | VI |

| | | | |
|---|---|---|---|
| Ga0196617_10000713 | CE10 | TISS1 | VI |
| Ga0196617_100007288 | GH3 | TISS1 | VI |
| Ga0196617_100007296 | CE4 | TISS1 | VI |
| Ga0196617_10000769 | CE1 | TISS1 | VI |
| Ga0196617_10001084 | AA3 | TISS1 | VI |
| Ga0196617_10001251 | CBM50 | TISS1 | VI |
| Ga0196617_10001251 | CBM50 | TISS1 | VI |
| Ga0196617_10001251 | CBM50 | TISS1 | VI |
| Ga0196617_10001251 | CBM50 | TISS1 | VI |
| Ga0196617_1000286 | CE10 | TISS1 | VI |
| Ga0196617_10003113 | GH18 | TISS1 | VI |
| Ga0196617_10004218 | CBM50 | TISS1 | VI |
| Ga0196617_10004240 | CE4 | TISS1 | VI |
| Ga0196617_10005416 | GH109 | TISS1 | VI |
| Ga0196617_10005467 | GH23 | TISS1 | VI |
| Ga0196617_1000051 | GH51 | CLOS1 | VI |
| Ga0196617_10000512 | GH29 | CLOS1 | VI |
| Ga0196617_100005134 | CE4 | CLOS1 | VI |
| Ga0196617_100005144 | GH43 | CLOS1 | VI |
| Ga0196617_10000517 | CE15 | CLOS1 | VI |
| Ga0196617_100005175 | GH31 | CLOS1 | VI |
| Ga0196617_10000518 | GH95 | CLOS1 | VI |
| Ga0196617_100005181 | CBM25 | CLOS1 | VI |
| Ga0196617_100005183 | GH9 | CLOS1 | VI |
| Ga0196617_100005190 | AA6 | CLOS1 | VI |
| Ga0196617_100005192 | GH51 | CLOS1 | VI |
| Ga0196617_100005195 | SLH | CLOS1 | VI |
| Ga0196617_100005195 | SLH | CLOS1 | VI |
| Ga0196617_100005195 | SLH | CLOS1 | VI |
| Ga0196617_10000520 | GH42 | CLOS1 | VI |
| Ga0196617_100005203 | GH2 | CLOS1 | VI |
| Ga0196617_10000521 | SLH | CLOS1 | VI |
| Ga0196617_100005216 | CBM50 | CLOS1 | VI |
| Ga0196617_100005269 | SLH | CLOS1 | VI |
| Ga0196617_100005269 | SLH | CLOS1 | VI |
| Ga0196617_1000053 | GH109 | CLOS1 | VI |
| Ga0196617_10000534 | GH35 | CLOS1 | VI |
| Ga0196617_10000548 | SLH | CLOS1 | VI |
| Ga0196617_10000548 | SLH | CLOS1 | VI |
| Ga0196617_10000548 | SLH | CLOS1 | VI |
| Ga0196617_1000055 | GH36 | CLOS1 | VI |
| Ga0196617_1000056 | GH36 | CLOS1 | VI |
| Ga0196617_10000567 | GH13 | CLOS1 | VI |
| Ga0196617_10000568 | GH36 | CLOS1 | VI |
| Ga0196617_1000057 | GH105 | CLOS1 | VI |
| Ga0196617_10000579 | GH10 | CLOS1 | VI |
| Ga0196617_1000058 | GH3 | CLOS1 | VI |
| Ga0196617_10000586 | GH43 | CLOS1 | VI |

| | | | |
|---|---|---|---|
| Ga0196617_10000587 | GH95 | CLOS1 | VI |
| Ga0196617_1000059 | GH31 | CLOS1 | VI |
| Ga0196617_100006113 | CE10 | CLOS1 | VI |
| Ga0196617_100006118 | GH43 | CLOS1 | VI |
| Ga0196617_100006212 | CE8 | CLOS1 | VI |
| Ga0196617_100006223 | CE3 | CLOS1 | VI |
| Ga0196617_100006228 | CBM22 | CLOS1 | VI |
| Ga0196617_100006228 | GH30 | CLOS1 | VI |
| Ga0196617_10000638 | GH18 | CLOS1 | VI |
| Ga0196617_10000669 | GH88 | CLOS1 | VI |
| Ga0196617_10000695 | GH109 | CLOS1 | VI |
| Ga0196617_100011115 | CBM50 | CLOS1 | VI |
| Ga0196617_100011143 | CE10 | CLOS1 | VI |
| Ga0196617_100011180 | GH3 | CLOS1 | VI |
| Ga0196617_10001122 | GH36 | CLOS1 | VI |
| Ga0196617_10001140 | SLH | CLOS1 | VI |
| Ga0196617_10001140 | SLH | CLOS1 | VI |
| Ga0196617_10001140 | SLH | CLOS1 | VI |
| Ga0196617_10001148 | GH109 | CLOS1 | VI |
| Ga0196617_100015121 | GH106 | CLOS1 | VI |
| Ga0196617_10001539 | GH67 | CLOS1 | VI |
| Ga0196617_10001577 | CE4 | CLOS1 | VI |
| Ga0196617_10001596 | CE15 | CLOS1 | VI |
| Ga0196617_10001596 | SLH | CLOS1 | VI |
| Ga0196617_10001596 | SLH | CLOS1 | VI |
| Ga0196617_10001596 | SLH | CLOS1 | VI |
| Ga0196617_100018109 | GH133 | CLOS1 | VI |
| Ga0196617_100018129 | GH2 | CLOS1 | VI |
| Ga0196617_10001814 | CBM50 | CLOS1 | VI |
| Ga0196617_10001830 | CE4 | CLOS1 | VI |
| Ga0196617_10001838 | CE4 | CLOS1 | VI |
| Ga0196617_10001876 | CE7 | CLOS1 | VI |
| Ga0196617_100020114 | GH112 | CLOS1 | VI |
| Ga0196617_100020115 | GH127 | CLOS1 | VI |
| Ga0196617_100020127 | GH63 | CLOS1 | VI |
| Ga0196617_10002017 | CBM50 | CLOS1 | VI |
| Ga0196617_1000203 | CBM16 | CLOS1 | VI |
| Ga0196617_10002031 | GH109 | CLOS1 | VI |
| Ga0196617_10002032 | CBM37 | CLOS1 | VI |
| Ga0196617_10002032 | SLH | CLOS1 | VI |
| Ga0196617_10002032 | SLH | CLOS1 | VI |
| Ga0196617_10002032 | SLH | CLOS1 | VI |
| Ga0196617_10002033 | CBM48 | CLOS1 | VI |
| Ga0196617_10002033 | CBM48 | CLOS1 | VI |
| Ga0196617_10002033 | GH13 | CLOS1 | VI |
| Ga0196617_10002040 | GH29 | CLOS1 | VI |
| Ga0196617_10002041 | GH95 | CLOS1 | VI |
| Ga0196617_10002055 | GH109 | CLOS1 | VI |

| | | | |
|---|---|---|---|
| Ga0196617_10002065 | GH43 | CLOS1 | VI |
| Ga0196617_10002092 | GH109 | CLOS1 | VI |
| Ga0196617_10002712 | CBM6 | CLOS1 | VI |
| Ga0196617_10002712 | GH2 | CLOS1 | VI |
| Ga0196617_1000274 | GH23 | CLOS1 | VI |
| Ga0196617_10002765 | GH109 | CLOS1 | VI |
| Ga0196617_10002766 | GH38 | CLOS1 | VI |
| Ga0196617_10002776 | GH105 | CLOS1 | VI |
| Ga0196617_10002777 | CE12 | CLOS1 | VI |
| Ga0196617_10002778 | GH105 | CLOS1 | VI |
| Ga0196617_10002780 | GH5 | CLOS1 | VI |
| Ga0196617_10002785 | CBM67 | CLOS1 | VI |
| Ga0196617_10002785 | GH78 | CLOS1 | VI |
| Ga0196617_1000344 | CE9 | CLOS1 | VI |
| Ga0196617_10003461 | GH2 | CLOS1 | VI |
| Ga0196617_10003466 | GH106 | CLOS1 | VI |
| Ga0196617_10003498 | GH38 | CLOS1 | VI |
| Ga0196617_10003636 | GH43 | CLOS1 | VI |
| Ga0196617_10003637 | GH43 | CLOS1 | VI |
| Ga0196617_10003643 | CE9 | CLOS1 | VI |
| Ga0196617_10003684 | CBM50 | CLOS1 | VI |
| Ga0196617_10003684 | CBM50 | CLOS1 | VI |
| Ga0196617_10003849 | GH15 | CLOS1 | VI |
| Ga0196617_10003870 | GH3 | CLOS1 | VI |
| Ga0196617_10003871 | SLH | CLOS1 | VI |
| Ga0196617_10003880 | GH2 | CLOS1 | VI |
| Ga0196617_10005513 | CE1 | CLOS1 | VI |
| Ga0196617_10005912 | GH28 | CLOS1 | VI |
| Ga0196617_10005913 | GH105 | CLOS1 | VI |
| Ga0196617_10005914 | GH43 | CLOS1 | VI |
| Ga0196617_10005958 | GH39 | CLOS1 | VI |
| Ga0196617_10005959 | GH95 | CLOS1 | VI |
| Ga0196617_10006041 | GH130 | CLOS1 | VI |
| Ga0196617_1000651 | CBM50 | CLOS1 | VI |
| Ga0196617_10006515 | PL9 | CLOS1 | VI |
| Ga0196617_10006557 | GH73 | CLOS1 | VI |
| Ga0196617_10006557 | SLH | CLOS1 | VI |
| Ga0196617_10006740 | PL12 | CLOS1 | VI |
| Ga0196617_10007036 | CBM50 | CLOS1 | VI |
| Ga0196617_10007036 | CBM50 | CLOS1 | VI |
| Ga0196617_10007036 | GH18 | CLOS1 | VI |
| Ga0196617_10007049 | GH94 | CLOS1 | VI |
| Ga0196617_1000758 | GH1 | CLOS1 | VI |
| Ga0196617_10007720 | GH109 | CLOS1 | VI |
| Ga0196617_10007733 | SLH | CLOS1 | VI |
| Ga0196617_10007733 | SLH | CLOS1 | VI |
| Ga0196617_10007748 | GH4 | CLOS1 | VI |
| Ga0196617_10007750 | CBM66 | CLOS1 | VI |

| | | | |
|---|---|---|---|
| Ga0196617_10007750 | PL11 | CLOS1 | VI |
| Ga0196617_10009210 | CE1 | CLOS1 | VI |
| Ga0196617_10009212 | GH13 | CLOS1 | VI |
| Ga0196617_10009249 | GH2 | CLOS1 | VI |
| Ga0196617_10010336 | GH106 | CLOS1 | VI |
| Ga0196617_10010340 | SLH | CLOS1 | VI |
| Ga0196617_1001112 | GH42 | CLOS1 | VI |
| Ga0196617_10011120 | GH3 | CLOS1 | VI |
| Ga0196617_10011122 | GH130 | CLOS1 | VI |
| Ga0196617_10011128 | GH88 | CLOS1 | VI |
| Ga0196617_10011135 | CBM67 | CLOS1 | VI |
| Ga0196617_10011135 | GH78 | CLOS1 | VI |
| Ga0196617_10011136 | GH3 | CLOS1 | VI |
| Ga0196617_10011620 | GH43 | CLOS1 | VI |
| Ga0196617_10011622 | CE10 | CLOS1 | VI |
| Ga0196617_10011624 | GH105 | CLOS1 | VI |
| Ga0196617_1001205 | GH4 | CLOS1 | VI |
| Ga0196617_10015018 | GH51 | CLOS1 | VI |
| Ga0196617_10015028 | SLH | CLOS1 | VI |
| Ga0196617_10015029 | CE4 | CLOS1 | VI |
| Ga0196617_1001561 | dockerin | CLOS1 | VI |
| Ga0196617_1001562 | SLH | CLOS1 | VI |
| Ga0196617_1001562 | SLH | CLOS1 | VI |
| Ga0196617_1001562 | SLH | CLOS1 | VI |
| Ga0196617_10015626 | GH4 | CLOS1 | VI |
| Ga0196617_1001563 | GH4 | CLOS1 | VI |
| Ga0196617_10016319 | CE7 | CLOS1 | VI |
| Ga0196617_10016826 | SLH | CLOS1 | VI |
| Ga0196617_1001793 | GH120 | CLOS1 | VI |
| Ga0196617_10019216 | CBM50 | CLOS1 | VI |
| Ga0196617_10019320 | CBM6 | CLOS1 | VI |
| Ga0196617_1002031 | GH109 | CLOS1 | VI |
| Ga0196617_1002039 | CBM54 | CLOS1 | VI |
| Ga0196617_1002039 | SLH | CLOS1 | VI |
| Ga0196617_1002039 | SLH | CLOS1 | VI |
| Ga0196617_1002039 | SLH | CLOS1 | VI |
| Ga0196617_10022112 | PL22 | CLOS1 | VI |
| Ga0196617_10022114 | GH18 | CLOS1 | VI |
| Ga0196617_10022114 | SLH | CLOS1 | VI |
| Ga0196617_10022114 | SLH | CLOS1 | VI |
| Ga0196617_1002911 | GH23 | CLOS1 | VI |
| Ga0196617_10029410 | GH10 | CLOS1 | VI |
| Ga0196617_1004705 | GH35 | CLOS1 | VI |
| Ga0196617_1005872 | SLH | CLOS1 | VI |
| Ga0187557_1002153 | CE9 | BWF2A | VI |
| Ga0187557_1002195 | CE4 | BWF2A | VI |
| Ga0187557_1002239 | SLH | BWF2A | VI |
| Ga0187557_1002239 | SLH | BWF2A | VI |

| | | | |
|---|---|---|---|
| Ga0187557_1002239 | SLH | BWF2A | VI |
| Ga0187557_100243 | GH57 | BWF2A | VI |
| Ga0187557_100286 | CE14 | BWF2A | VI |
| Ga0187557_1003196 | CE1 | BWF2A | VI |
| Ga0187557_1003298 | GH109 | BWF2A | VI |
| Ga0187557_100357 | SLH | BWF2A | VI |
| Ga0187557_100357 | SLH | BWF2A | VI |
| Ga0187557_1004109 | GH23 | BWF2A | VI |
| Ga0187557_100472 | GH13 | BWF2A | VI |
| Ga0187557_100494 | CE4 | BWF2A | VI |
| Ga0187557_10055 | SLH | BWF2A | VI |
| Ga0187564_10115 | SLH | SW3C | VI |
| Ga0187564_10115 | SLH | SW3C | VI |
| Ga0187564_10115 | SLH | SW3C | VI |
| Ga0187564_10665 | SLH | SW3C | VI |
| Ga0187564_10665 | SLH | SW3C | VI |
| Ga0187564_10665 | SLH | SW3C | VI |
| Ga0187564_1086 | SLH | SW3C | VI |
| Ga0187557_100244+Ga0187564_103146 | GH57 | SW3C + BWF2A | VI |
| Ga0187564_101173+Ga0187557_1001187 | CE1 | SW3C + BWF2A | VI |
| Ga0187564_10433+Ga0187557_100434 | GH23 | SW3C + BWF2A | VI |
| Ga0187557_1003195+Ga0196617_1003937+Ga0187564_102192 | CE10 | SW3C + BWF2A + COPR1 | VI |
| Ga0187564_10247+Ga0196617_1005403+Ga0187557_100347 | GH74 | SW3C + BWF2A + COPR1 | VI |
| Ga0187564_10674+Ga0187557_1002248+Ga0196617_1003705 | GH16 | SW3C + BWF2A + Unbinned | VI |
| Ga0187564_10675+Ga0187557_1002249+Ga0196617_1003704 | GH3 | SW3C + BWF2A + Unbinned | VI |
| Ga0196617_10009442+Ga0187564_10367 | CBM66 | SW3C + COPR1 | VI |
| Ga0187564_10357+Ga0196617_1014501 | CE4 | SW3C + Unbinned | VI |
| Ga0196617_10006346 | AA6 | RCLO1 | VII |
| Ga0196617_10021811 | CE8 | RCLO1 | VII |
| Ga0196617_100004229 | CBM50 | TISS1 | VII |
| Ga0196617_10001282 | AA3 | TISS1 | VII |
| Ga0196617_10002881 | CE4 | TISS1 | VII |
| Ga0196617_10002771 | GH105 | CLOS1 | VII |
| Ga0196617_10019218 | GH18 | CLOS1 | VII |
| Ga0196617_1008651 | CBM6 | RCLO1 | VIII |
| Ga0196617_1008651 | GH2 | RCLO1 | VIII |
| Ga0196617_1006112 | CE4 | COPR1 | VIII |
| Ga0196617_1008452 | CE4 | COPR1 | VIII |
| Ga0196617_100006226 | SLH | CLOS1 | VIII |
| Ga0196617_100006226 | SLH | CLOS1 | VIII |
| Ga0196617_100006226 | SLH | CLOS1 | VIII |
| Ga0196617_100006227 | GH10 | CLOS1 | VIII |
| Ga0196617_10001149 | GH31 | CLOS1 | VIII |
| Ga0196617_1005975 | CBM35 | CLOS1 | VIII |
| Ga0196617_1005975 | GH26 | CLOS1 | VIII |
| Ga0196617_1007811 | CBM32 | CLOS1 | VIII |
| Ga0196617_1007811 | PL12 | CLOS1 | VIII |
| Ga0196617_1008533 | GH130 | CLOS1 | VIII |

| | | | |
|---|---|---|---|
| Ga0196617_1003511 | GH109 | COPR2 | VIII |
| Ga0196617_1002713 | PL12 | COPR3 | VIII |
| Ga0196617_1002715 | PL12 | COPR3 | VIII |
| Ga0187557_100130 | SLH | BWF2A | VIII |
| Ga0187557_100130 | SLH | BWF2A | VIII |
| Ga0187557_100130 | SLH | BWF2A | VIII |
| Ga0187557_1002251 | CBM35 | BWF2A | n.a. |
| Ga0187557_1002251 | GH18 | BWF2A | n.a. |
| Ga0187557_100245 | GH57 | BWF2A | n.a. |
| Ga0187557_10084 | GH8 | BWF2A | n.a. |
| Ga0187557_10091 | Dockerin | BWF2A | n.a. |
| Ga0187557_10091 | Dockerin | BWF2A | n.a. |
| Ga0187557_10091 | GH9 | BWF2A | n.a. |
| Ga0187557_10201 | CBM3 | BWF2A | n.a. |
| Ga0187557_10212 | Dockerin | BWF2A | n.a. |
| Ga0187557_10212 | GH8 | BWF2A | n.a. |
| Ga0196617_10006232+Ga0187564_103145 | GH57 | SW3C + COPR1 | n.a. |
| Ga0187564_102275 | GH73 | SW3C | n.a. |
| Ga0187564_10616 | CE4 | SW3C | n.a. |
| Ga0187564_1133 | GH9 | SW3C | n.a. |
| Ga0187564_1292 | CBM3 | SW3C | n.a. |

**Supplementary Table 7. Genome IMG ID and SRA numbers for datasets used in this study**

| Dataset | SRA Accession number | IMG Genome ID |
|---|---|---|
| **Metagenomes** | | |
| SEM1b_D1B | SRX3777359 | 3300019285 |
| SEM1b_D2B | SRX3777360 | 3300019285 |
| SEM1b_D1B-r1 (16S rRNA) | SRX3777358 | |
| SEM1b_D2B-r1 (16S rRNA) | SRX3777363 | |
| SEM1b_D1B-r2 (16S rRNA) | SRX3777361 | |
| SEM1b_D2B-r2 (16S rRNA) | SRX3777362 | |
| **Isolate genomes** | | |
| *C. proteolyticus BWF2A* | SRX3783215 | 2731957509 |
| *C. proteolyticus SW3C* | SRX3783214 | 2731957514 |
| **SEM1b time series** | | |
| **Dataset** | **Metatranscriptomes** | **16S rRNA** |
| SEM1b_T1A | | SRX3777365 |
| SEM1b_T1B | | SRX3777364 |
| SEM1b_T1C | | SRX3777357 |
| SEM1b_T2A | SRX3777356 | SRX3777346 |
| SEM1b_T2B | SRX3777345 | SRX3777348 |
| SEM1b_T2C | SRX3777347 | SRX3777350 |
| SEM1b_T3A | SRX3777349 | SRX3777352 |
| SEM1b_T3B | SRX3777351 | SRX3777354 |
| SEM1b_T3C | SRX3777353 | SRX3777368 |
| SEM1b_T4A | SRX3777369 | SRX3777370 |
| SEM1b_T4B | SRX3777371 | SRX3777372 |
| SEM1b_T4C | SRX3777373 | SRX3777374 |
| SEM1b_T5A | SRX3777375 | SRX3777366 |
| SEM1b_T5B | SRX3777367 | SRX3777344 |
| SEM1b_T5C | SRX3777343 | SRX3777342 |
| SEM1b_T6A | SRX3777341 | SRX3777340 |
| SEM1b_T6B | SRX3777339 | SRX3777338 |
| SEM1b_T6C | SRX3777337 | SRX3777336 |
| SEM1b_T7A | SRX3777335 | SRX3777384 |
| SEM1b_T7B | SRX3777385 | SRX3777382 |
| SEM1b_T7C | SRX3777383 | SRX3777380 |
| SEM1b_T8A | SRX3777381 | SRX3777378 |
| SEM1b_T8B | SRX3777379 | SRX3777376 |
| SEM1b_T8C | SRX3777377 | SRX3777355 |

**REFERENCES.**


Azizi A, Kim W, Lee JH (2016). Comparison of microbial communities during the anaerobic digestion of Gracilaria under mesophilic and thermophilic conditions. *World Journal of Microbiology and Biotechnology* **32:** 1-17.

Chen Y, Zhang F, Wang T, Shen N, Yu ZW, Zeng RJ (2016). Hydraulic retention time affects stable acetate production from tofu processing wastewater in extreme-thermophilic (70 °C) mixed culture fermentation. *Bioresource Technology* **216:** 722-728.

Chen Y, Xiao K, Jiang X, Shen N, Zeng RJ, Zhou Y (2017). In-situ sludge pretreatment in a single-stage anaerobic digester. *Bioresource Technology* **238:** 102-108.

Fitamo T, Treu L, Boldrin A, Sartori C, Angelidaki I, Scheutz C (2017). Microbial population dynamics in urban organic waste anaerobic co-digestion with mixed sludge during a change in feedstock composition and different hydraulic retention times. *Water Research* **118:** 261-271.

Gagliano MC, Braguglia CM, Gianico A, Mininni G, Nakamura K, Rossetti S (2015). Thermophilic anaerobic digestion of thermal pretreated sludge: Role of microbial community structure and correlation with process performances. *Water Research* **68:** 498-509.

Hagen LH, Frank JA, Zamanzadeh M, Eijsink VGH, Pope PB, Horn SJ *et al* (2017). Quantitative metaproteomics highlight the metabolic contributions of uncultured phylotypes in a thermophilic anaerobic digester. *Applied and Environmental Microbiology* **83**.

Ho D, Jensen P, Gutierrez-Zamora ML, Beckmann S, Manefield M, Batstone D (2016). High-rate, high temperature acetotrophic methanogenesis governed by a three population consortium in anaerobic bioreactors. *PLoS ONE* **11:** 1-13.

Kobayashi T, Li YY, Harada H (2008). Analysis of microbial community structure and diversity in the thermophilic anaerobic digestion of waste activated sludge. *Water Science and Technology* **57:** 1199-1205.

Kouzuma A, Tsutsumi M, Ishii Si, Ueno Y, Abe T, Watanabe K (2017). Non-autotrophic methanogens dominate in anaerobic digesters. *Scientific Reports* **7:** 1-13.

Lü F, Bize A, Guillot A, Monnet V, Madigou C, Chapleur O *et al* (2014). Metaproteomics of cellulose methanisation under thermophilic conditions reveals a surprisingly high proteolytic activity. *ISME Journal* **8:** 88-102.

Pervin HM, Dennis PG, Lim HJ, Tyson GW, Batstone DJ, Bond PL (2013). Drivers of microbial community composition in mesophilic and thermophilic temperature-phased anaerobic digestion pre-treatment reactors. *Water Research* **47:** 7098-7108.

Sun W, Yu G, Louie T, Liu T, Zhu C, Xue G *et al* (2015). From mesophilic to thermophilic digestion: the transitions of anaerobic bacterial, archaeal, and fungal community structures in sludge and manure samples. *Applied Microbiology and Biotechnology* **99:** 10271-10282.

Yu B, Lou Z, Zhang D, Shan A, Yuan H, Zhu N *et al* (2015). Variations of organic matters and microbial community in thermophilic anaerobic digestion of waste activated sludge with the addition of ferric salts. *Bioresource Technology* **179:** 291-298.
